# The adoption of a westernized gut microbiome in Indian Immigrants and Indo-Canadians is associated with dietary acculturation

**DOI:** 10.1101/2024.03.04.582285

**Authors:** Leah D. D’Aloisio, Mamatha Ballal, Sanjoy Ghosh, Natasha Haskey, Nijiati Abulizi, Ramin Karimianghadim, Chuyi Liu, Pacha Sruthi, Lakshmipriya Nagarajan, Sudha Vasudevan, Vignesh Shetty, Mrudgandha Purandare, Ushasi Bhaumik, Debaki Ranjan Howlader, Sepideh Pakpour, Jacqueline Barnett, Deanna L. Gibson

## Abstract

Indian immigration to westernized countries has recently surged, increasing their risk of Inflammatory Bowel Disease (IBD) post-migration. While crucial for understanding IBD risk, the gut microbiome remains understudied in Indians. This cross-sectional study examines the gut microbiomes of Indians residing in India, Indo-Immigrants, and Indo-Canadians in comparison to Euro-Canadian and Euro-Immigrant controls to understand the impact of westernization on their gut. Stool samples for 16S rRNA and shotgun sequencing assessed microbial taxa and functional profiles, complemented by dietary and demographic data to evaluate lifestyle patterns. Results revealed distinct microbiotas in Indians and Indo-Immigrants compared to control groups, with high abundances of *Prevotella* spp. and CAZymes reflecting their high complex carbohydrate diet. Indo-Canadians exhibited a transitional microbiome towards westernization, mirroring increasing dietary acculturation. Considering 44% of Canadians are first- and second-generation immigrants and the global adoption of westernized practices, future research should investigate the health implications of such microbiome transitions in immigrant populations and newly industrialized nations.

## BACKGROUND

The growing immigrant population in North America is rapidly westernizing through the adoption of practices like reduced physical activity, changes in hygiene/medical practices, and acculturation of the westernized diet, commonly characterized by caloric dense, highly processed foods with low fiber intake. Amidst this backdrop are health disparities emerging in these populations. Furthermore, the global spread of industrialization raises concern for the increasing prevalence of modern diseases.^1^ For instance, inflammatory bowel disease (IBD) is increasing in prevalence globally, and alarming evidence reveals a specifically increased risk in young Indian immigrants and Indians born in several westernized countries.^2–4^ In 2019, India had an incidence rate of 2.34, whereas South Asians in Canada showed an incidence rate of 14.6 per 100,000 person-years, which was comparable to the general population in Canada.^5,6^ A likely contributor to this elevated risk in westernized countries like Canada may stem from changes in the gut microbiome as a response to living in an industrialized environment.

The industrialized microbiome has been previously characterized by the decline of VANISH (volatile and/or associated negatively with industrialized societies of humans) taxa, frequently found in populations adhering to traditional lifestyles, and the rise of BloSSUM (bloom or selected in societies of urbanization/modernization) taxa, which appear in modern, industrialized individuals.^7,8^ Furthermore, health consequences following the assimilation into westernized culture have been previously documented. For example, non-industrialized Irish Travellers who adopted modern practices and Hmong and Karen migrants from Thailand who relocated to the United States (US) both experienced gut microbiome shifts that were associated with an increased risk for metabolic disorders.^9,10^ Currently, there is only one study published on the Indian immigrant gut microbiome, highlighting the gap in non-western/non-European microbiome research. Indeed, over 71% of all microbiome research comes from westernized cohorts, skewing our interpretation of a healthy human gut.^11^ Here, we aimed to investigate the distinction between the Indian and westernized gut microbiomes and the differences in the gut of Indians who migrate to Canada, alongside the adoption of the westernized diet.

## METHODS

### Study Design

A cross-sectional study was conducted including three Indian groups (Indian ancestry was confirmed in each group): (1) Indians residing in India (“Indians”), (2) Indians who migrated to Canada (“Indo-Immigrants”) and (3) Canadians born in Canada from Indian ancestry (“Indo-Canadians”). Caucasian individuals born in Canada with European ancestry (“Euro-Canadian”) were used as a control, and immigrants with European ancestry from a westernized country (“Euro-Immigrants”) as a westernized immigrant control. Healthy participants aged 17-55 years were recruited. Exclusion criteria included pregnancy, diagnosis of any chronic inflammatory condition, antibiotics used or travel to India within three months prior to stool collection. Participants completed a demographic questionnaire to assess baseline characteristics. Immigrants also completed a lifestyle survey to assess changes in lifestyle and immigration-related stress.

### Stool Collection & DNA Extraction

Stool samples were collected from India (Kolkata and Manipal) and Canada (Kelowna) and processed on site as previously described.^12^ DNA was sent to Gut4Health for 16S rRNA gene sequencing on the Illumina MiSeq platform (V4-V4, 515F/806R primers) (∼75,800 reads per sample) and to the [Center for Health Genomics and Informatics](https://cumming.ucalgary.ca/research/cat/health-genomics/home) for shotgun sequencing on the Illumina NovaSeq platform (∼9.9M reads per sample). Additional details are in Supplementary Methods.

### Taxonomic Analysis

Demultiplexed 16S rRNA gene reads were processed in QIIME 2 (Version 2023.9)^13^ with DADA2^14^ for quality control and taxonomic classification with GreenGenes2 (10.28.22).^15^ Demultiplexed raw shotgun reads underwent quality assessment with FastQC/MultiQC and used KneadData (Version 0.10) for trimming and quality control. Taxonomic profiling was conducted using MetaPhlAn4 for higher taxonomic resolution at the species level,^16^ and each sample was normalized to relative abundances. Diversity metrics included Shannon and Pielou’s Evenness (Kruskal-Wallis), and Bray Curtis and Weighted UniFrac (PERMANOVA). Differentially abundant bacteria were determined using the Linear Discriminate Analysis (LDA) Effect Size (LEfSe) algorithm with a 3.5 threshold and alpha 0.05.^17^ Using shotgun data, the normalized relative abundance MetaPhlAn4 output table was imported into MicrobiomeAnalyst 2.0 to apply a Random Forest algorithm ranked according to their Mean Decrease Accuracy score, indicating the extent to which each taxon’s presence or absence influences the overall accuracy of classifying samples into their respective cohort.^18^ These taxa were then visualized in a heatmap. Using 16S rRNA gene data, BugBase was used to predict microbiome phenotypes,^19,20^ and associations between beta diversity, dietary patterns and baseline characteristics were analyzed with distance-based redundancy analysis (dbDRA). Additional details for taxonomic analysis methods can be found in Supplementary Methods.

### Functional Profiling

Shotgun reads were analyzed for functional potential using HUMAnN (Version 3.6).^21^ Gene family and pathway abundances were annotated using the UniRef90^22^ and MetaCyc^23^ databases, respectively, then normalized to reads per kilobase to account for gene length. Both tables were further normalized to relative abundance to account for sampling depth. The normalized gene family abundance output was then regrouped to the CAZy and KEGG databases. To determine differentially abundant genes/pathways across cohorts, tables were collapsed to perform LEfSe analysis on the unstratified data using a one-against-all strategy with a threshold of 3.0 on the logarithmic LDA score and an alpha of 0.05 for Kruskal-Wallis test among classes.^17^ Differentially abundant features were then plotted as a heatmap using MicrobiomeAnalyst 2.0.^18^

### Nutritional Analysis

Participants filled out a 3-day food log prior to their stool collection, then researchers retrospectively inputted this data into the ESHA Food Processor® software (Version 11.11.0) to obtain the calculated nutrient intake for each subject. To mitigate bias from using a North American nutritional software, Indian and Indo-Immigrant food logs were also entered into the MDRF EpiNu® Nutritional software, tailored for Indian populations. Additional details on our approach for dietary analysis can be found in Supplementary Methods.

### Statistical Analysis

Statistical analyses for demographics and dietary data were performed using both R statistical software and GraphPad Prism (Version 10.0.3). We applied the Kruskal-Wallis test coupled with Dunn’s post hoc analysis for nonparametric data. For data that passed the Shapiro-Wilk test of normality, an ordinary one-way ANOVA followed by Tukey’s post hoc test was performed. Categorical data were analyzed using either a Chi-Square, Fisher’s Exact or Fisher’s Exact with Monte Carlo simulation, depending on the data structure. Results were reported as median values and interquartile range (IQR), unless otherwise specified.

## RESULTS

### Participant Characteristics

The study included Indians (*n* = 61), Indo-Immigrants (*n* = 32), Indo-Canadians (*n*= 17), Euro-Canadians (*n* = 41) and Euro-Immigrants (*n* = 23) (Figure S1). Baseline demographics were comparable across groups, though Indians were older and consumed significantly less alcohol (p_BONF_ = 3.29e-09) (Table 1). Indo-Immigrants exhibited more of a change towards westernization (p = 0.002) and higher immigration-related stress (p = 0.04), with no significant differences in age at or years since immigration (Table 2).

**Table 1.**
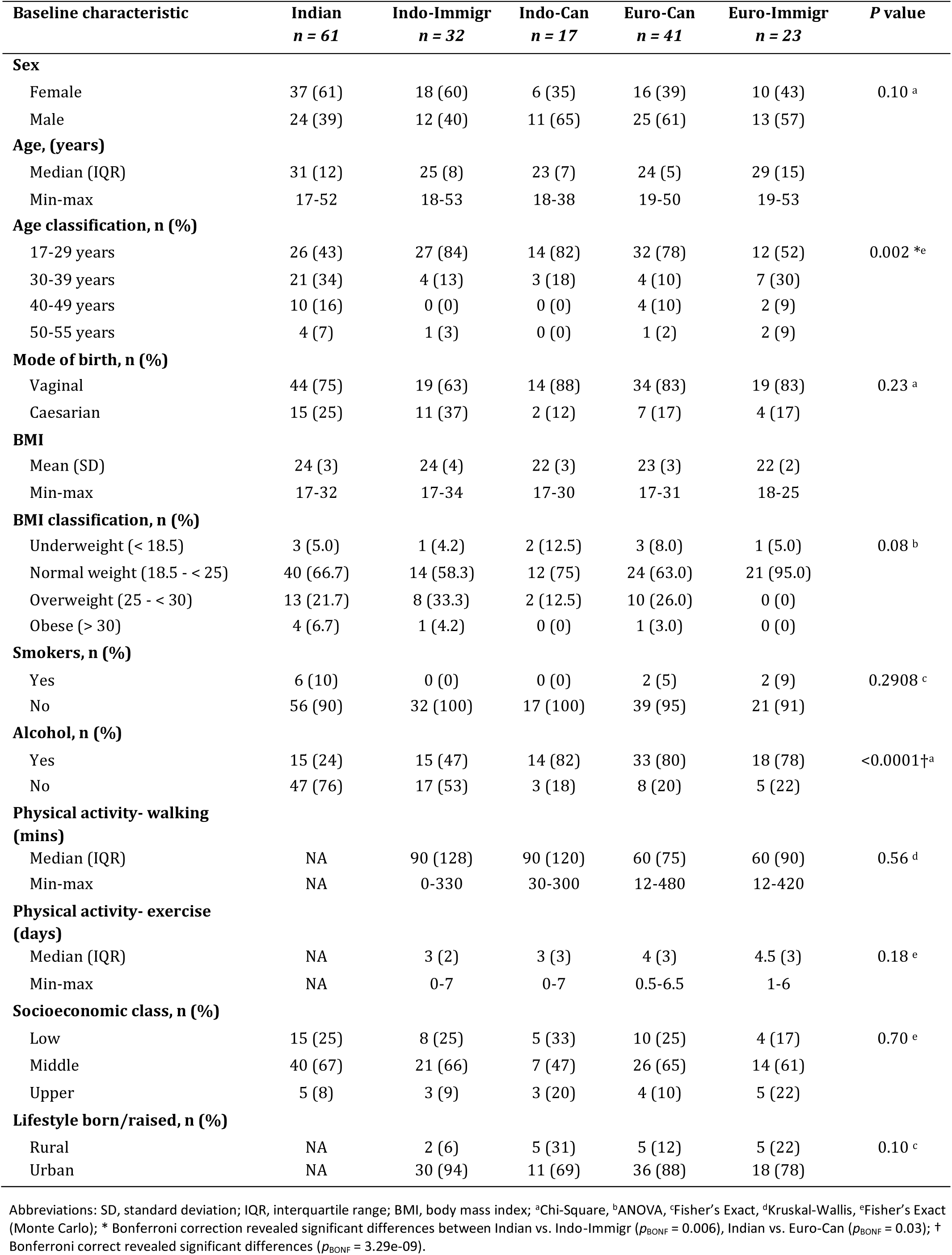
Baseline Characteristics of Participants.

| Baseline characteristic | Indian<br>*n* = 61 | Indo-Immigr<br>*n* = 32 | Indo-Can<br>*n* = 17 | Euro-Can<br>*n* = 41 | Euro-Immigr<br>*n* = 23 | *P* value |
| --- | --- | --- | --- | --- | --- | --- |
| **Sex** |  |  |  |  |  |  |
| Female | 37 (61) | 18 (60) | 6 (35) | 16 (39) | 10 (43) | 0.10 <sup>a</sup> |
| Male | 24 (39) | 12 (40) | 11 (65) | 25 (61) | 13 (57) |  |
| **Age, (years)** |  |  |  |  |  |  |
| Median (IQR) | 31 (12) | 25 (8) | 23 (7) | 24 (5) | 29 (15) |  |
| Min-max | 17-52 | 18-53 | 18-38 | 19-50 | 19-53 |  |
| **Age classification, n (%)** |  |  |  |  |  |  |
| 17-29 years | 26 (43) | 27 (84) | 14 (82) | 32 (78) | 12 (52) | 0.002 <sup>*e</sup> |
| 30-39 years | 21 (34) | 4 (13) | 3 (18) | 4 (10) | 7 (30) |  |
| 40-49 years | 10 (16) | 0 (0) | 0 (0) | 4 (10) | 2 (9) |  |
| 50-55 years | 4 (7) | 1 (3) | 0 (0) | 1 (2) | 2 (9) |  |
| **Mode of birth, n (%)** |  |  |  |  |  |  |
| Vaginal | 44 (75) | 19 (63) | 14 (88) | 34 (83) | 19 (83) | 0.23 <sup>a</sup> |
| Caesarian | 15 (25) | 11 (37) | 2 (12) | 7 (17) | 4 (17) |  |
| **BMI** |  |  |  |  |  |  |
| Mean (SD) | 24 (3) | 24 (4) | 22 (3) | 23 (3) | 22 (2) |  |
| Min-max | 17-32 | 17-34 | 17-30 | 17-31 | 18-25 |  |
| **BMI classification, n (%)** |  |  |  |  |  |  |
| Underweight (< 18.5) | 3 (5.0) | 1 (4.2) | 2 (12.5) | 3 (8.0) | 1 (5.0) | 0.08 <sup>b</sup> |
| Normal weight (18.5 - < 25) | 40 (66.7) | 14 (58.3) | 12 (75) | 24 (63.0) | 21 (95.0) |  |
| Overweight (25 - < 30) | 13 (21.7) | 8 (33.3) | 2 (12.5) | 10 (26.0) | 0 (0) |  |
| Obese (> 30) | 4 (6.7) | 1 (4.2) | 0 (0) | 1 (3.0) | 0 (0) |  |
| **Smokers, n (%)** |  |  |  |  |  |  |
| Yes | 6 (10) | 0 (0) | 0 (0) | 2 (5) | 2 (9) | 0.2908 <sup>c</sup> |
| No | 56 (90) | 32 (100) | 17 (100) | 39 (95) | 21 (91) |  |
| **Alcohol, n (%)** |  |  |  |  |  |  |
| Yes | 15 (24) | 15 (47) | 14 (82) | 33 (80) | 18 (78) | <0.0001<sup>†a</sup> |
| No | 47 (76) | 17 (53) | 3 (18) | 8 (20) | 5 (22) |  |
| **Physical activity- walking (mins)** |  |  |  |  |  |  |
| Median (IQR) | NA | 90 (128) | 90 (120) | 60 (75) | 60 (90) | 0.56 <sup>d</sup> |
| Min-max | NA | 0-330 | 30-300 | 12-480 | 12-420 |  |
| **Physical activity- exercise (days)** |  |  |  |  |  |  |
| Median (IQR) | NA | 3 (2) | 3 (3) | 4 (3) | 4.5 (3) | 0.18 <sup>e</sup> |
| Min-max | NA | 0-7 | 0-7 | 0.5-6.5 | 1-6 |  |
| **Socioeconomic class, n (%)** |  |  |  |  |  |  |
| Low | 15 (25) | 8 (25) | 5 (33) | 10 (25) | 4 (17) | 0.70 <sup>e</sup> |
| Middle | 40 (67) | 21 (66) | 7 (47) | 26 (65) | 14 (61) |  |
| Upper | 5 (8) | 3 (9) | 3 (20) | 4 (10) | 5 (22) |  |
| **Lifestyle born/raised, n (%)** |  |  |  |  |  |  |
| Rural | NA | 2 (6) | 5 (31) | 5 (12) | 5 (22) | 0.10 <sup>c</sup> |
| Urban | NA | 30 (94) | 11 (69) | 36 (88) | 18 (78) |  |

Abbreviations: SD, standard deviation; IQR, interquartile range; BMI, body mass index; <sup>a</sup>Chi-Square, <sup>b</sup>ANOVA, <sup>c</sup>Fisher's Exact, <sup>d</sup>Kruskal-Wallis, <sup>e</sup>Fisher's Exact (Monte Carlo); \* Bonferroni correction revealed significant differences between Indian vs. Indo-Immigr ( $p_{\text{BONF}} = 0.006$ ), Indian vs. Euro-Can ( $p_{\text{BONF}} = 0.03$ ); <sup>†</sup> Bonferroni correct revealed significant differences ( $p_{\text{BONF}} = 3.29\text{e-}09$ ).

**Table 2.** Characteristics of Immigrant Participants.

| Characteristic | Indo-Immigr<br>*n* = 32 | Euro-Immigr<br>*n* = 23 | *P* value |
| --- | --- | --- | --- |
| **Δ westernization, n (%)** |  |  |  |
| No westernization | 0 (0) | 4 (18) | 0.002 <sup>a</sup> |
| Little westernization | 12 (38) | 13 (59) |  |
| Moderate westernization | 17 (53) | 4 (18) |  |
| High westernization | 3 (9) | 1 (4) |  |
| **Immigration-related stress, n (%)** |  |  |  |
| Low stress | 22 (69) | 21 (91) | 0.04 <sup>a</sup> |
| Moderate stress | 8 (25) | 2 (9) |  |
| High stress | 2 (6) | 0 (0) |  |
| **Age at immigration, n (%)** |  |  |  |
| 0-3 years | 0 (0) | 1 (4) | 0.19 <sup>b</sup> |
| 4-17 years | 9 (28) | 3 (13) |  |
| 18-49 years | 23 (72) | 19 (83) |  |
| **Years since immigration** |  |  |  |
| Median (IQR) | 4 (6) | 3 (9) | 0.65 <sup>c</sup> |
| Min-max | 0-22 | 0-48 |  |

Abbreviations: IQR, interquartile range; <sup>a</sup>Chi-Square for Trend, <sup>b</sup>Fisher's Exact, <sup>c</sup>Mann-Whitney

### Indian Microbiota is Distinctive from Westernized Groups

To examine the taxa and complexity of the microbial ecosystem in Indians compared to westerners, alpha diversity revealed the lowest diversity in Indians and highest in Euro-Canadians (Figure 1A-B). Average percentage of unclassified reads per sample in each cohort can be found in Table S1. Bray Curtis dissimilarity of shotgun sequence data showed significant differences across groups, with 26.3% of variation explained on the first 2 axes (Figure 1C). Indians and Indo-Immigrants’ microbiotas differed from westernized cohorts (Table 3), and each other (pseudo-F = 9.132, p_BONF_ = 0.01). Weighted UniFrac also demonstrated distinctions in the Indian and Indo-Immigrant microbiomes (pseudo-F = 25.98, p_BONF_ = 0.01), with 61.8% of the variation explained on the first 2 axes (Figure 1D) with similar findings from 16S rRNA gene amplicon sequence data (Figure S2).

**Figure 1.**
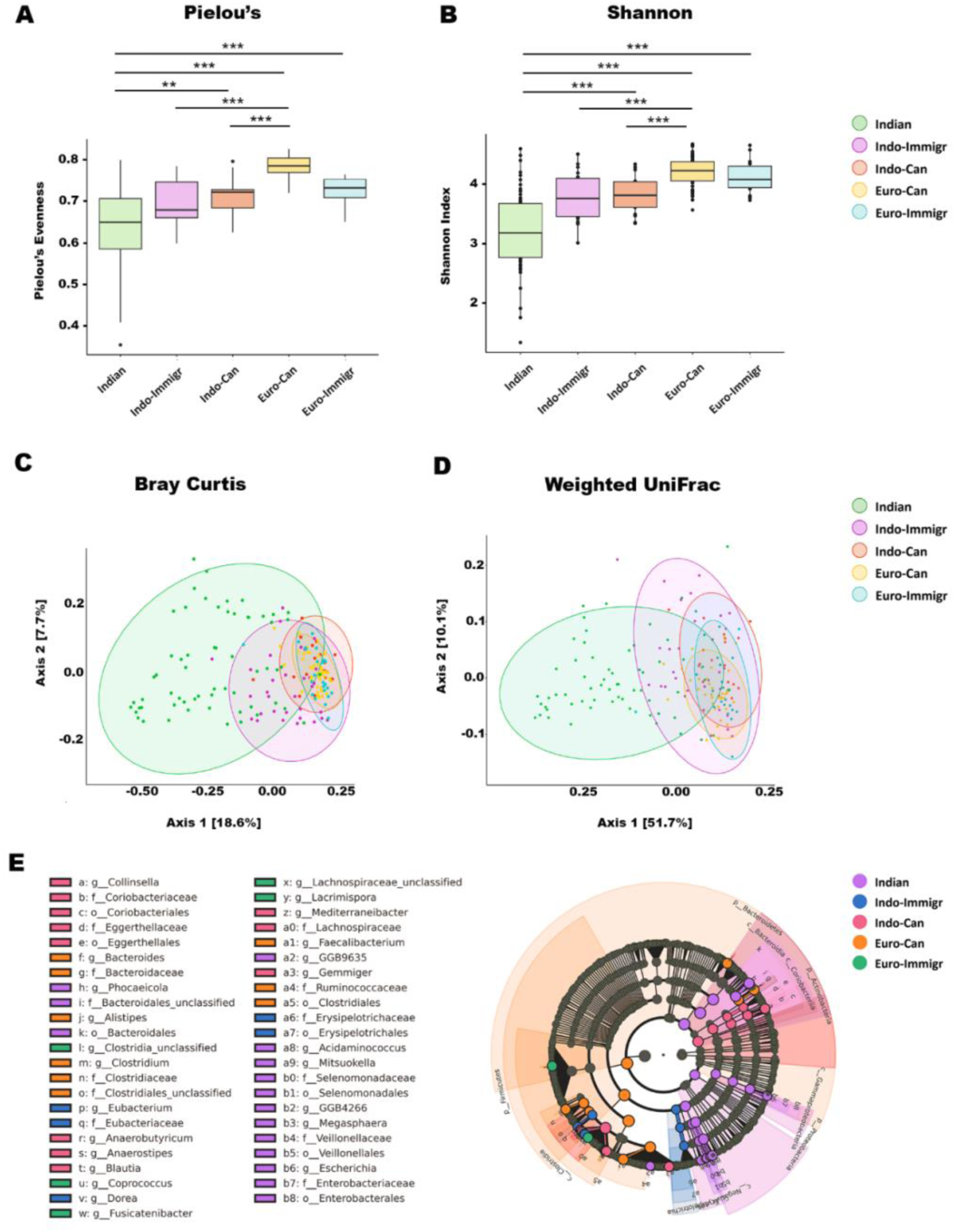
Indian and Indo-Immigrant Microbiotas are Distinct from Westernized Groups. **(A)** Pielou’s Evenness (H = 85.7, p = 1.07e-17). **(B)** Shannon’s Diversity (H = 79.8, p = 1.89e-16). Beta diversity was explored with Bray Curtis dissimilarity and Weighted UniFrac, using Pairwise Permutational Multivariate Analysis of Variance (PERMANOVA) to test differences between groups. **(C)** Bray Curtis principal coordinate analysis (PCoA) plot shows 26.3% of variation was captured on the first two axes. **(D)** Weighted UniFrac PCoA plot shows 61.8% of the variation was captured on the first two axes. Pairwise comparisons results can be found in Table S1. **(E)** LEfSe (Linear discriminate analysis Effect Size) cladogram results, depicting differentially abundant bacteria across cohorts, with genus set as lowest taxonomic rank. *** = q ≤ 0.001, ** = q ≤ 0.01, * = q ≤ 0.05. Boxes represent the interquartile range (IQR) between the first and third quartiles, the horizontal line indicates the median, and whiskers are the upper and lower values within 1.5 times the IQR.

**Table 3.** PERMANOVA Results from Beta Diversity Analysis.

| Pair | Bray Curtis |  | Weighted UniFrac |  |
| --- | --- | --- | --- | --- |
|  | pseudo-F | P value | pseudo-F | P value |
| Indo-Immigr vs. Indo-Can | 4.215003 | 0.01 | 4.670629 | 0.01 |
| Indo-Immigr vs. Euro-Can | 8.976951 | 0.01 | 11.932016 | 0.01 |
| Indo-Immigr vs. Euro-Immigr | 5.313757 | 0.01 | 6.903393 | 0.01 |
| Indo-Immigr vs. Indian | 9.132465 | 0.01 | 25.984365 | 0.01 |
| Indo-Can vs. Euro-Can | 3.084292 | 0.01 | 5.409267 | 0.01 |
| Indo-Can vs. Euro-Immigr | 2.166793 | 0.01 | 2.360739 | 0.10 |
| Indo-Can vs. Indian | 8.660676 | 0.01 | 25.206253 | 0.01 |
| Euro-Can vs. Euro-Immigr | 1.163757 | 1.00 | 1.116192 | 1.00 |
| Euro-Can vs. Indian | 18.284343 | 0.01 | 49.611215 | 0.01 |
| Euro-Immigr vs. Indian | 11.182855 | 0.01 | 31.565399 | 0.01 |

Adjusted *P* values are displayed, which were obtained using 999 permutations and Bonferroni correction applied. Beta diversity data from shotgun sequence data.

LEfSe identified dominant taxa in each group, with Bacteroidota (formerly Bacteroidetes) and Pseudomonadota (formerly Proteobacteria) in Indians and Firmicutes in Euro-Canadians (Figure 1E, Table S1). Indo-Immigrants displayed highest abundances of *Eubacterium* and Erysipelotrichaceae (a VANISH taxon)^24^, both previously found to be enriched in non-industrialized populations^25^, though abundance plots of both taxa reveal low abundances in Indians (Figure S4). As expected, genera common in the industrialized microbiome were enriched in the westernized European cohorts such as *Alistipes*, *Bacteroides*, and Clostridia, all previously noted at high abundances in industrialized groups.^25,26^ Nonetheless, average relative abundances for *Alistipes* and *Bacteroides* were higher in Indians than Indo-Immigrants. Indo-Canadians represented a transitional gut microbiota, harbouring both taxa found in non-industrialized, such as *Collinsella*, and industrialized populations, such as *Blautia* and *Anaerostipes* (both noted as BloSSUM taxa).^25,26^ Similar results were found in 16S rRNA gene sequence data (Figure S3, Table S2).

When assessing differences in bacterial species abundances amongst groups, LEfSe showed that Indians harboured higher abundances of *Phocaeicola plebeius*, *Megasphaera sp.*, *Dialister hominis,* and *Escherichia coli* (Figure 2). Among Indo-Immigrants, *Dorea longicatena*, *Faecalibacterium prausnitizii* and *Lachnospiraceae bacterium* were notably abundant, while Indo-Canadians had increased abundances of two *Ruminoccocus torques* SGBs, *Blautia wexlerae, Eubacterium rectale, Blautia massiliensis,* and *Anaerostipes hadrus*. Taxa enriched in Euro-Canadians included *Phocaeicola vulgatus*, *Lachnospiraceae bacterium, Clostridium sp*., and *Blautia faecis*, and Euro-Immigrants showed enrichment of *Fusicatenibacter saccharivorans, Clostridia bacterium,* and *Blautia obeum.* A key distinguishing feature from Indians to Indo-Canadians was the decline in *Prevotella* spp., with *Prevotella copri* (clade A) being over five times more abundant in Indians (Table 4). *P. copri* Clade A relative abundances were 10.8 for Indians, 5.89 for Indo-Immigrants, and 2.75 for Indo-Canadians (Figure 2C). A similar pattern of decreasing abundance across Indian cohorts was also observed for Clades B, C and D. All clades were lowest in abundance in Euro-Canadians. Overall, these findings demonstrate that the gut microbiome in Indians are distinct from the West, and Indo-Canadians exhibit a transition towards the industrialized microbiome.

**Figure 2.**
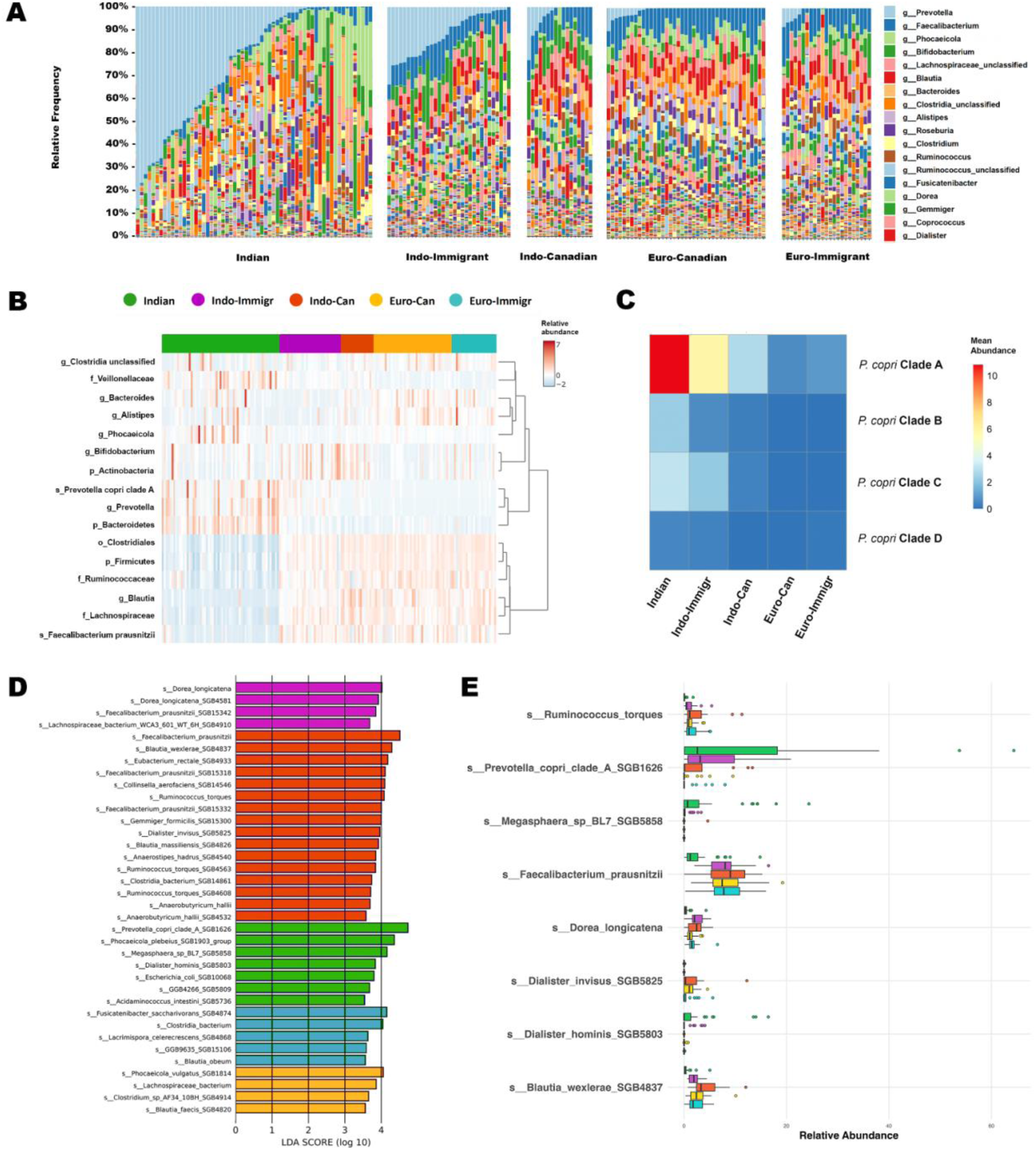
Significantly Different Gut Bacterial Abundances Detected Across Cohorts. **(A)** Taxonomic stacked bar plots depicting relative abundances of top 18 most abundant bacteria across all samples from shotgun sequence data (N = 174; Indian (n = 61), Indo-Immigr (n = 32), Indo-Can (n = 23), Euro-Can (n = 41), Euro-Immigr (n = 23). **(B)** Heatmap generated in MicrobiomeAnalyst 2.0, displaying taxa identified by Random Forest as key features that contribute to the predictive accuracy of classifying samples into their respective groups (Figure S5). For the heatmap, features were filtered for minimum 4 counts in 20% or more samples, and low variance filter of 10%. Bars on the top represent the cohorts that group the individual sample columns displayed. Taxa are labelled in rows, with taxonomic rank noted before the bacteria name. Colours on heatmap represent the relative abundances of each bacteria in a given sample. **(C)** Heatmap generated in R Studio, displaying the average relative abundances of *P. copri* clades in each cohort. **(D)** LEfSe results of top differentially abundant bacteria. A Kruskal-Wallis test was performed with a significance (α) at 0.05 for one-against-all comparisons. Differentially abundant bacteria were detected using a Linear Discriminate Analysis (LDA) score (equal to or greater than 3.5). **(E)** Abundance plot of notable differentially abundant species identified by LEfSe, displaying average relative abundance per cohort.

**Table 4.**
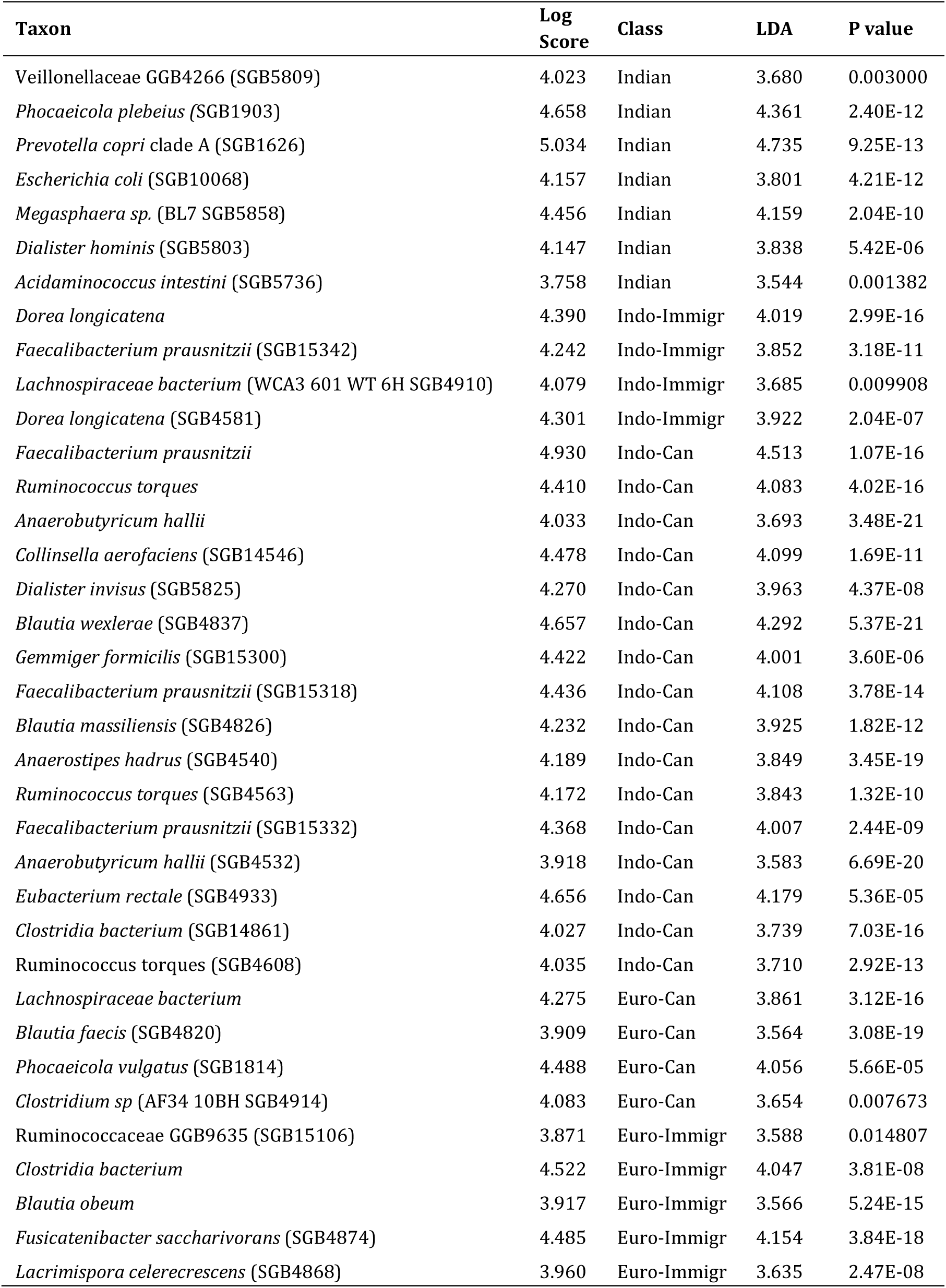
Differentially abundant bacteria from shotgun sequence data.

### Indian Gut Microbiome Predicted to be More Pathogen-Tolerant

To understand potential functionality of the gut microbiota in Indians, we used BugBase predictions (Figure 3), which estimated that Indians appeared to demonstrate higher stress tolerance driven by Pseudomonadota (Proteobacteria). The Indian cohorts had the highest potentially pathogenic bacteria (p_FDR_ = 2.07e-18) mostly dominated by Gram negative bacteria driven mainly by a combination of Pseudomonadota, Bacteroidota (Bacteroidetes), and Firmicutes. In addition, BugBase predicted that the Indian gut microbiome had the highest potential for biofilm formation. Overall, these results suggest the Indian gut microbiome may be more pathogen-tolerant, as healthy individuals were predicted to harbour microbes with higher pathogenic potential and stress tolerance. However, it is important to note that results from BugBase are only predictions made using an older GreenGenes database, therefore these findings should be further substantiated in future studies.

**Figure 3.**
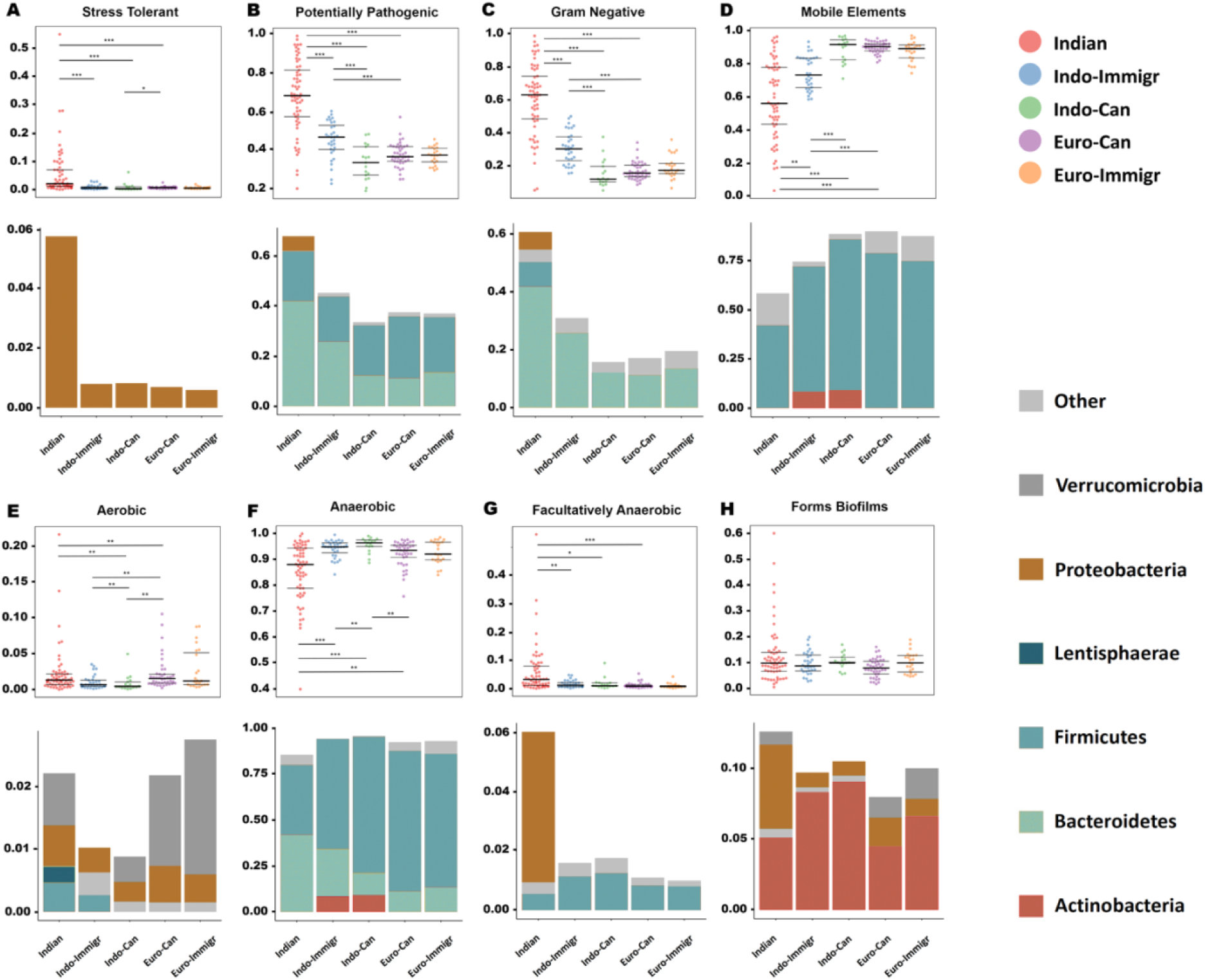
BugBase Predicts Higher Pathogenic Potential and Stress-Tolerant Microbiome in Indians. A Kruskal-Wallis test was performed in BugBase followed by Mann-Whitney-Wilcoxon tests for pairwise comparisons with adjusted *P* values shown. Relative abundance is presented on the y-axis. Indian (n = 61), Indo-Immigr (n = 32), Indo-Can (n = 23), Euro-Can (n = 41), Euro-Immigr (n = 23). **(A)** Stress tolerance **(B)** Potentially pathogenic bacteria **(C)** Gram negative bacteria **(D)** Containing mobile elements **(E)** Aerobic bacteria **(F)** Anaerobic bacteria **(G)** Facultatively Anaerobic bacteria **(H)** Biofilm formation. Taxonomic contributions are also displayed for each phenotype prediction. *** = p ≤ 0.001, ** = p ≤ 0.01, * = p ≤ 0.05

### Gut Functional Potential in Indians is Distinctive from Westernized Groups

To further understand functional attributes of the Indian gut microbiome, pathway analysis using MetaCyc was used and revealed pathways enriched in lipopolysaccharide (LPS) production, peptidoglycan synthesis, microbial growth/metabolism, protein synthesis and DNA synthesis/repair (Figure 4). In Indo-Canadians, indications of increased glucose storage were detected. Euro-Canadians had increased pathways related to fatty acid biosynthesis, and Euro-Immigrants expressed pathways indicating sugar metabolism (Table S3). CAZy gene family analysis revealed distinct profiles in the Indian gut including glycoside hydrolases (GH)43, GH51, and GH10, all of which are known for their degradation of plant-based polysaccharides (Table S4). *P. copri* was the top taxa contributing to these CAZy families (Figure 5A). In addition, KEGG identities K00561 and K18220 were enriched in Indians and Indo-Immigrants, respectively, both recognized for antimicrobial resistance genes (Figure 5B). Overall, these results suggest that both environmental factors, diets and drugs, were influencing the gut microbiome of Indians.

**Figure 4.**
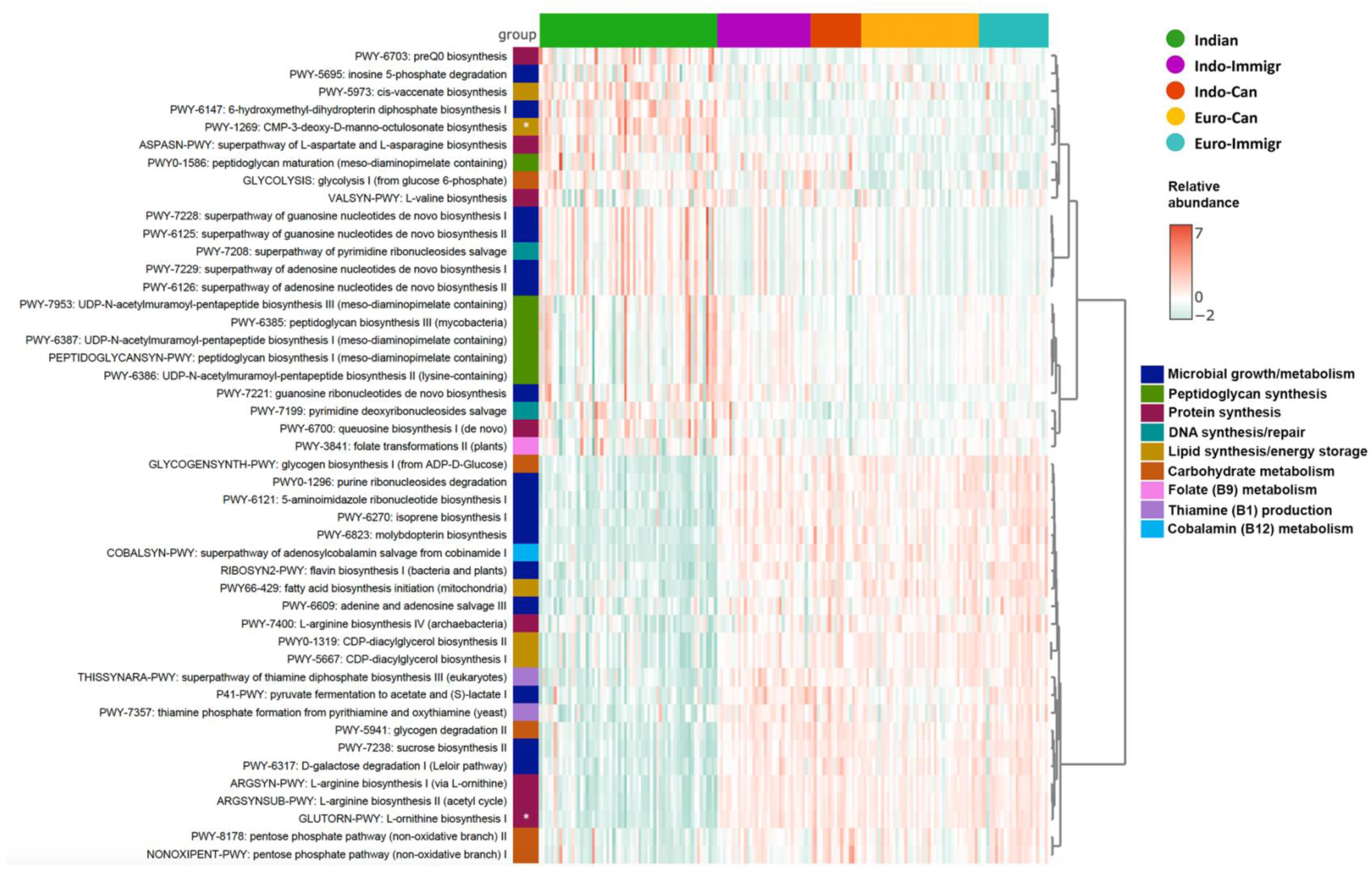
Microbial Metabolic Pathways Show Distinction in Functional Potential of Indian Gut Microbiome. Pathway abundances were annotated using MetaCyc, then normalized to account for gene length and sampling depth. LEfSe (Linear discriminate analysis Effect Size) was used to test for differentially abundant microbial metabolic pathways using unstratified data. A Kruskal-Wallis test was performed with a significance (a) at 0.05 for one-against-all comparisons. Discriminate features were identified using a Linear Discriminate Analysis (LDA) score, with an LDA score equal to or greater than 3.0. Using MicrobiomeAnalyst 2.0, a heatmap was generated to display the differentially abundant pathways identified from LEfSe. Each column is a sample, and the coloured bars on top represent the cohorts. Indian (n = 61), Indo-Immigr (n = 32), Indo-Can (n = 23), Euro-Can (n = 41), Euro-Immigr (n = 23). Metabolic pathways are labelled in rows and annotated with a colour that describes their general function. Colours on the heatmap represent the relative abundances of each pathway. * *L-orthinine biosynthesis I* is indirectly linked with protein synthesis and also involved with urea cycle; * *CMP-3-deoxy-D-manno-octulosonate biosynthesis* is directly involved in lipopolysaccharide (LPS) production.

**Figure 5.**
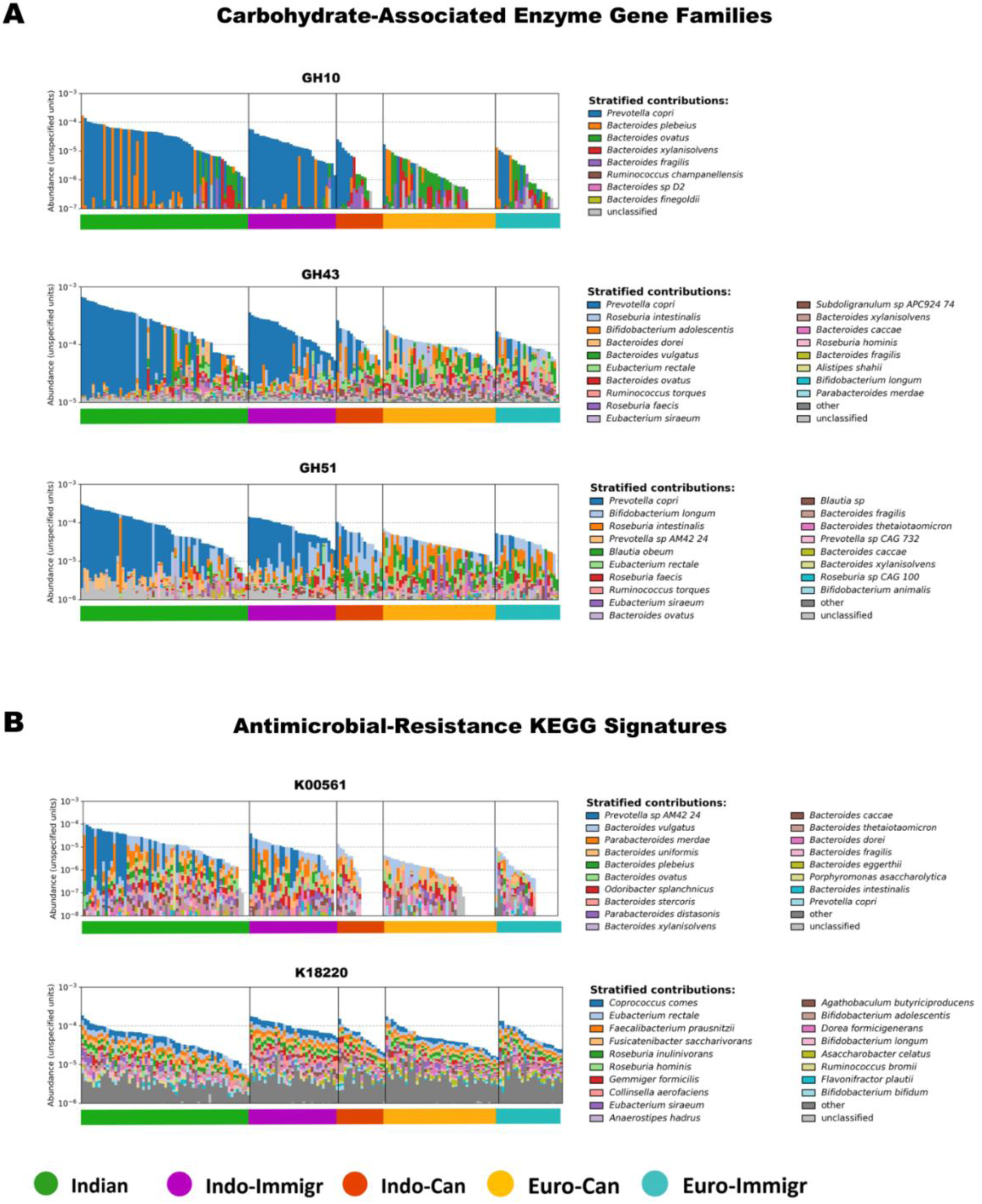
*Prevotella copri* is the Top Taxa Contributing to CAZy Families Abundant in Indians. Normalized gene family abundance table was regrouped to CAZy database using *human_regroup_table*. Barplots for stratified taxa contributions for each CAZy gene family was generated using *human_barplot*. Indian (n = 61), Indo-Immigr (n = 32), Indo-Can (n = 23), Euro-Can (n = 41), Euro-Immigr (n = 23). *Prevotella copri* is the top contributor to CAZyme families **(A)** GH10, GH43, GH51, all which were enriched in Indians. **(B)** Top taxa contributions to antimicrobial resistance-related KEGG orthologues abundant in Indians and Indo-Immigrants. Normalized gene family abundance table was regrouped to KEGG database using *human_regroup_table*. Barplots for stratified taxa contributions for each KEGG ortholog was generated using *human_barplot*. *Prevotella copri AM42 24* is the top contributor to K00561, enriched in Indians and *Coprococcus comes* is the top contributing taxa to K18220, enriched in Indo-Immigrants.

### Westernized Dietary Acculturation Observed in Indo-Canadians

To understand the role of diet associated with the gut microbiome in our Indian cohort, we collected and analyzed dietary recalls using both validated software from North America and from India to account for Indian foods. Indo-Canadians had the highest daily intake of NOVA Group 4 Ultra-Processed Food (UPF), constituting 61% of their caloric intake (95% CI = 467.9 – 780.5 kcal, per 1000 kcal), while Indians consumed significantly less than all other cohorts, reporting 12% of their daily intake (95% CI = 63.66 – 127.1 kcal per 1000 kcal) (Figure 6). Consistent with high UPF intake, total fiber was also found to be the lowest in Indo-Canadians, opposite in Indians who consumed more fiber than Indo-Canadians (p = 0.0028) and Euro-Canadians (p = 0.0110). The highest proportion of non-meat eaters were Indians at 60% (*n* = 36/61, p = <0.0001), with 30% vegetarian and 30% pescetarian. In Indo-Immigrants, 19% (*n* = 6/32), while 12% (*n* = 2/17) of Indo-Canadians, 10% (*n* = 4/41) Euro-Canadians and 39% (*n* = 9/23) of Euro-Immigrants were non-meat eaters. Cooking oil usage highlighted sunflower oil, ghee and mustard oil as the top choices of Indians, whereas Indo-Immigrants preferred olive oil, butter and ghee, with olive oil and butter being the primary fats used by the three westernized groups (Figure S6, Table 5). These results highlight the westernized diet adopted by Indo-Canadians, including high levels of processed food and low levels of fiber, are potential drivers in the westernized gut microbiota.

**Figure 6.**
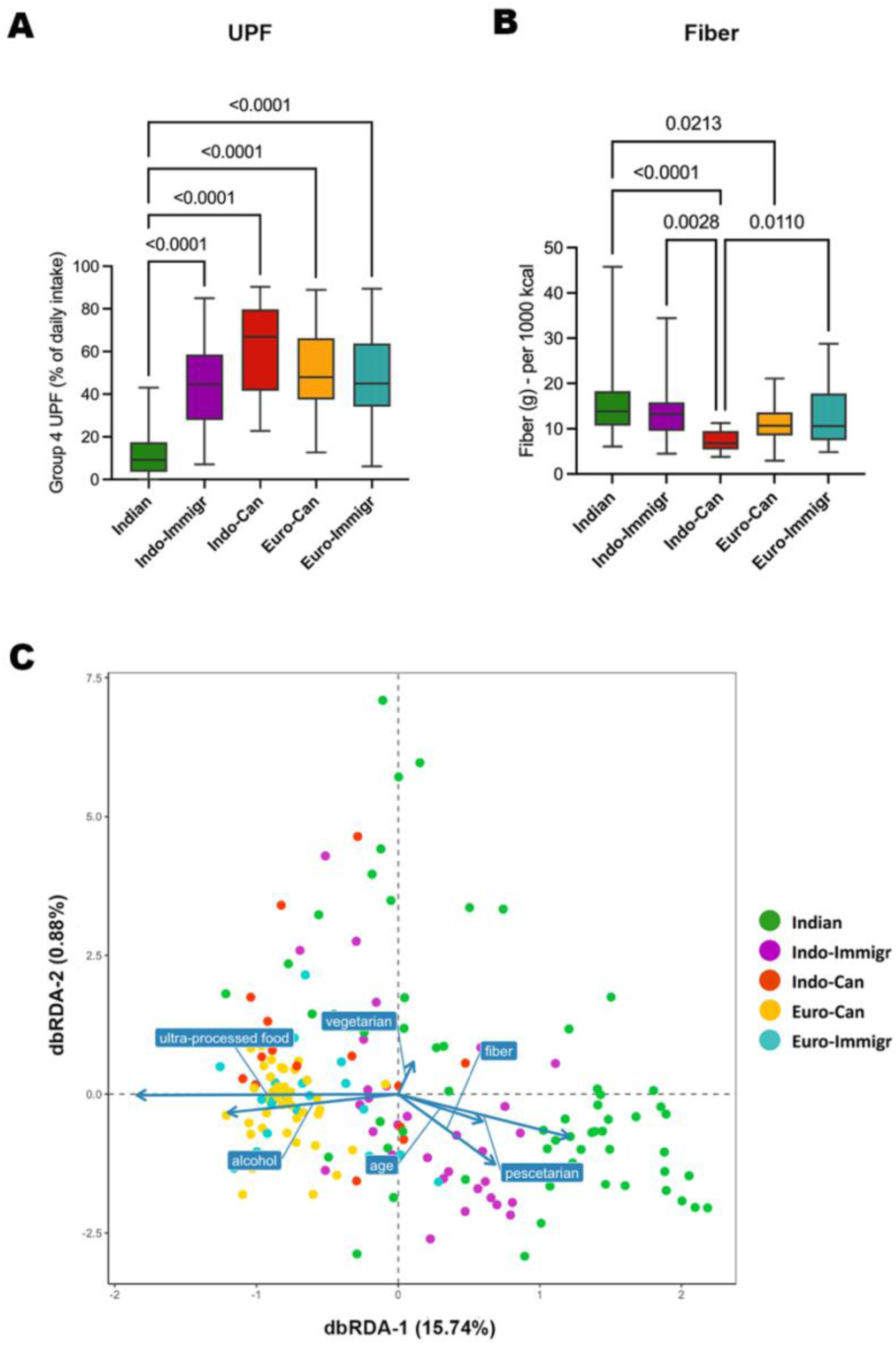
Significant Intake of Ultra-Processed Food and Alcohol are Drivers in the Westernized Gut Microbiota. Middle bands are the median values, top and bottom boxes display the first and third quartiles, and whiskers are the min and max values. Indian (n = 61), Indo-Immigr (n = 32), Indo-Can (n = 23), Euro-Can (n = 41), Euro-Immigr (n = 23). **(A)** Calories from Group 4 Ultra-Processed Food (UPF) was counted for each subject from the ESHA dietary reports then compared to total average daily caloric intake. A Kruskal-Wallis test was performed, followed by Dunn’s multiple comparisons, with adjusted p values displayed. **(B)** Fiber intake was normalized per 1000 kcal for each subject. An ordinary one-way ANOVA was performed, followed by Tukey’s multiple comparisons, with adjusted p values displayed. **(C)** Distance-based redundancy analysis (dbRDA) plot was generated using Weighted UniFrac distance matrix from 16S rRNA gene amplicon data. This dbRDA plot depicts the distribution of samples and their associations with lifestyle factors. Each dot is a sample, colour coded by cohort. Variance inflation factor values ranged between 1.14 (vegetarian) – 1.38 (pescetarian), indicating each variable uniquely contributed to the model.

**Table 5.** Percentages of respondents who reported use of each cooking oil.

| **Oil Type** | **Indian**<br>**$n = 43$** | **Indo-Immigr**<br>**$n = 18$** | **Indo-Can**<br>**$n = 9$** | **Euro-Can**<br>**$n = 35$** | **Euro-Immigr**<br>**$n = 18$** |
| --- | --- | --- | --- | --- | --- |
| Butter | 5% | 24% | 27% | 24% | 26% |
| Ghee | 12% | 16% | 0% | 0% | 0% |
| Coconut | 9% | 4% | 0% | 5% | 10% |
| Olive | 1% | 28% | 45% | 44% | 39% |
| Sunflower | 30% | 0% | 0% | 2% | 0% |
| Canola | 0% | 12% | 0% | 13% | 6% |
| Vegetable | 5% | 4% | 18% | 2% | 10% |
| Mustard | 11% | 8% | 0% | 0% | 0% |
| Avocado | 0% | 0% | 0% | 5% | 6% |
| Groundnut | 8% | 0% | 0% | 0% | 0% |
| Rice | 8% | 0% | 0% | 0% | 0% |
| Other | 9% | 4% | 9% | 4% | 3% |

To understand what dietary factors could be driving microbial taxa, the dbRDA model, accounting for demographics and dietary patterns elucidated 16.62% of the microbial variance on the first 2 axes (PERMANOVA F = 5.9923, p = 0.001) (Figure 6C). After false discovery rate (FDR) adjustment, UPF and alcohol intake were principal determinants in differentiating the microbiota in westernized cohorts (p_FDR_ = 0.003). The pescetarian diet (p_FDR_ = 0.070) and fiber intake (p_FDR_ = 0.2268) appeared to influence clustering of Indian samples, albeit not significant. Advancing age (p_FDR_ = 0.044) influenced another clustering, whereas vegetarianism showed no significant effect (p_FDR_ = 0.371). Overall, these results suggest that UPF and alcohol intake were driving westernization of the gut microbiome in Indian migrants with this effect most prominent in Indo-Canadians. Furthermore, the higher fiber intake observed in Indians correlated with high *Prevotella* spp. and CAZy gene families tailored to the more traditional Indian diet.

Given the above data indicated that the Indo-Canadian group had adopted a westernized diet, further analysis of diet was examined including absolute macronutrient and micronutrient intakes (Table 6-7; Table S5-S6). Total energy intake was significantly higher in Euro-Canadian males vs. Indo-Immigrant males (p = 0.0144, 95% CI = −1461 – −110.0 kcal). Median percentage of protein intake was highest in Euro-Canadians (16.1%, 95% CI = 13.2 – 17.5%) and lowest protein intake was in Indians (12.5%, 95% CI = 11.7 – 13.3). Indians consumed the highest percentage of carbohydrates in their diet (56.5%, 95% CI = 54.7% - 59.0%) and lowest carbohydrate diet intake was in Indo-Canadians (44.0%, 95% CI = 40.6% – 50.6%). The highest percentage of fat consumption was in Indo-Canadians (39.7%, CI 95% 33.8% – 42.2%) and the lowest was in Indians (31.1%, 95% CI = 28.5% – 34.4%). Software comparisons between North American ESHA and Indian EpiNu revealed no macronutrient composition differences however, EpiNu calculated a significantly higher intake of PUFAs and MUFAs and lower intake of SFA amongst Indians, highlighting discrepancy in dietary fat types between dietary software, which could skew dietary analysis between the countries (Fig 7A-B). When the type of oils consumed were examined, it was revealed that Indo-Canadians have the least diversity with no traditional fats added to their diet like ghee (Fig 7C).

**Table 6.**
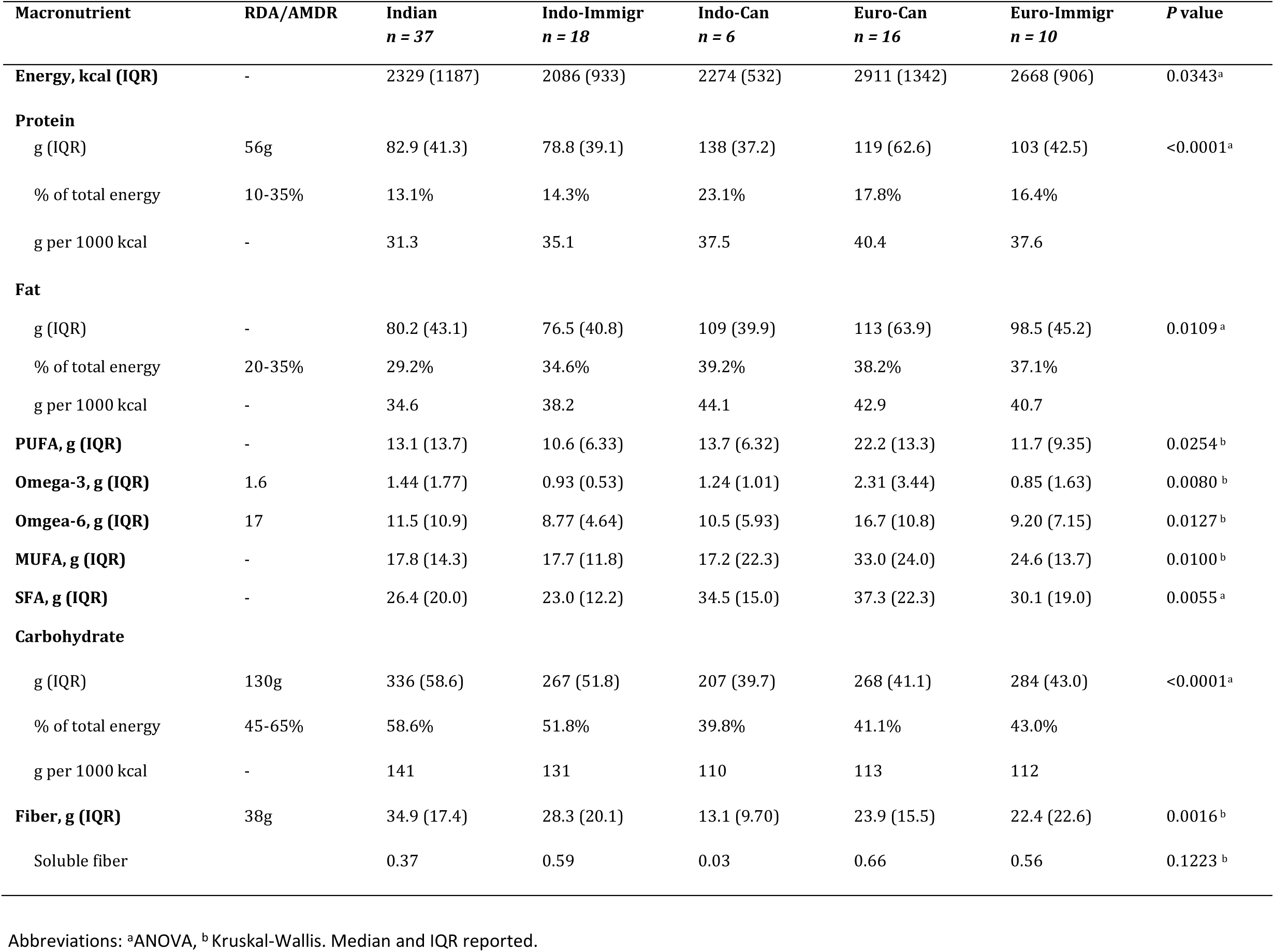
Male participant absolute macronutrient intake.

| Macronutrient | RDA/AMDR | Indian<br>*n* = 37 | Indo-Immigr<br>*n* = 18 | Indo-Can<br>*n* = 6 | Euro-Can<br>*n* = 16 | Euro-Immigr<br>*n* = 10 | *P* value |
| --- | --- | --- | --- | --- | --- | --- | --- |
| **Energy, kcal (IQR)** | - | 2329 (1187) | 2086 (933) | 2274 (532) | 2911 (1342) | 2668 (906) | 0.0343<sup>a</sup> |
| **Protein** |  |  |  |  |  |  |  |
| g (IQR) | 56g | 82.9 (41.3) | 78.8 (39.1) | 138 (37.2) | 119 (62.6) | 103 (42.5) | <0.0001<sup>a</sup> |
| % of total energy | 10-35% | 13.1% | 14.3% | 23.1% | 17.8% | 16.4% |  |
| g per 1000 kcal | - | 31.3 | 35.1 | 37.5 | 40.4 | 37.6 |  |
| **Fat** |  |  |  |  |  |  |  |
| g (IQR) | - | 80.2 (43.1) | 76.5 (40.8) | 109 (39.9) | 113 (63.9) | 98.5 (45.2) | 0.0109<sup>a</sup> |
| % of total energy | 20-35% | 29.2% | 34.6% | 39.2% | 38.2% | 37.1% |  |
| g per 1000 kcal | - | 34.6 | 38.2 | 44.1 | 42.9 | 40.7 |  |
| **PUFA, g (IQR)** | - | 13.1 (13.7) | 10.6 (6.33) | 13.7 (6.32) | 22.2 (13.3) | 11.7 (9.35) | 0.0254<sup>b</sup> |
| **Omega-3, g (IQR)** | 1.6 | 1.44 (1.77) | 0.93 (0.53) | 1.24 (1.01) | 2.31 (3.44) | 0.85 (1.63) | 0.0080<sup>b</sup> |
| **Omega-6, g (IQR)** | 17 | 11.5 (10.9) | 8.77 (4.64) | 10.5 (5.93) | 16.7 (10.8) | 9.20 (7.15) | 0.0127<sup>b</sup> |
| **MUFA, g (IQR)** | - | 17.8 (14.3) | 17.7 (11.8) | 17.2 (22.3) | 33.0 (24.0) | 24.6 (13.7) | 0.0100<sup>b</sup> |
| **SFA, g (IQR)** | - | 26.4 (20.0) | 23.0 (12.2) | 34.5 (15.0) | 37.3 (22.3) | 30.1 (19.0) | 0.0055<sup>a</sup> |
| **Carbohydrate** |  |  |  |  |  |  |  |
| g (IQR) | 130g | 336 (58.6) | 267 (51.8) | 207 (39.7) | 268 (41.1) | 284 (43.0) | <0.0001<sup>a</sup> |
| % of total energy | 45-65% | 58.6% | 51.8% | 39.8% | 41.1% | 43.0% |  |
| g per 1000 kcal | - | 141 | 131 | 110 | 113 | 112 |  |
| **Fiber, g (IQR)** | 38g | 34.9 (17.4) | 28.3 (20.1) | 13.1 (9.70) | 23.9 (15.5) | 22.4 (22.6) | 0.0016<sup>b</sup> |
| Soluble fiber |  | 0.37 | 0.59 | 0.03 | 0.66 | 0.56 | 0.1223<sup>b</sup> |

Abbreviations: <sup>a</sup>ANOVA, <sup>b</sup>Kruskal-Wallis. Median and IQR reported.

**Table 7.** Female participant absolute macronutrient intake.

| Macronutrient | RDA/AMDR | Indian<br>*n* = 24 | Indo-Immigr<br>*n* = 14 | Indo-Can<br>*n* = 11 | Euro-Can<br>*n* = 25 | Euro-Immigr<br>*n* = 13 | *P* value |
| --- | --- | --- | --- | --- | --- | --- | --- |
| **Energy, kcal (IQR)** | - | 1747 (630) | 1976 (1136) | 1812 (527) | 2265 (910) | 2227 (1060) | 0.0332* |
| **Protein** |  |  |  |  |  |  |  |
| g (IQR) | 46g | 51.5 (21.8) | 65.3 (47.9) | 63.1 (20.5) | 81.0 (20.8) | 66.6 (50.7) | 0.0004<sup>b</sup> |
| % of total energy | 10-35% | 12.3% | 13.8% | 12.9% | 14.1% | 14.0% |  |
| g per 1000 kcal | - | 31.3 | 35.1 | 37.5 | 40.4 | 37.6 |  |
| **Fat** |  |  |  |  |  |  |  |
| g (IQR) | - | 62.7 (32.3) | 69.8 (38.0) | 73.1 (27.5) | 89.2 (40.4) | 103 (57.0) | 0.0018<sup>b</sup> |
| % of total energy | 20-35% | 33.4% | 34.4% | 39.7% | 38.7% | 36.7% |  |
| g per 1000 kcal | - | 34.6 | 38.2 | 44.1 | 42.9 | 40.7 |  |
| **PUFA, g (IQR)** | - | 12.1 (9.49) | 12.9 (11.0) | 8.06 (4.33) | 13.3 (8.90) | 9.87 (13.0) | ns |
| **Omega-3, g (IQR)** | 1.1 | 1.24 (0.85) | 1.46 (1.02) | 0.84 (0.44) | 1.42 (1.25) | 1.06 (1.16) | ns |
| **Omega-6, g (IQR)** | 12 | 10.5 (8.72) | 8.80 (8.25) | 7.00 (4.04) | 10.0 (8.92) | 8.73 (7.67) | ns |
| **MUFA, g (IQR)** | - | 16.5 (10.5) | 15.6 (13.4) | 15.3 (9.77) | 23.8 (13.7) | 19.0 (23.6) | ns |
| **SFA, g (IQR)** | - | 18.3 (14.9) | 23.4 (10.6) | 22.9 (5.66) | 30.6 (14.3) | 41.7 (35.1) | ns |
| **Carbohydrate** |  |  |  |  |  |  |  |
| g (IQR) | 130g | 258 (86.4) | 256 (116) | 218 (99.5) | 272 (119) | 248 (141) | ns |
| % of total energy | 45-65% | 55.7% | 53.6% | 38.4% | 47.6% | 49.4% |  |
| g per 1000 kcal | - | 141 | 131 | 110 | 113 | 112 |  |
| **Fiber, g (IQR)** | 25g | 24.6 (17.8) | 23.1 (18.7) | 14.7 (6.30) | 27.0 (15.2) | 28.8 (28.4) | 0.0163<sup>a</sup> |
| Soluble fiber |  | 0.21 | 0.69 | 0.05 | 0.75 | 0.45 | 0.0003<sup>b</sup> |

Abbreviations: <sup>a</sup>ANOVA, <sup>b</sup>Kruskal-Wallis. Median and IQR reported.
\*Tukey's post hoc analysis revealed no significant differences between groups

**Figure 7.**
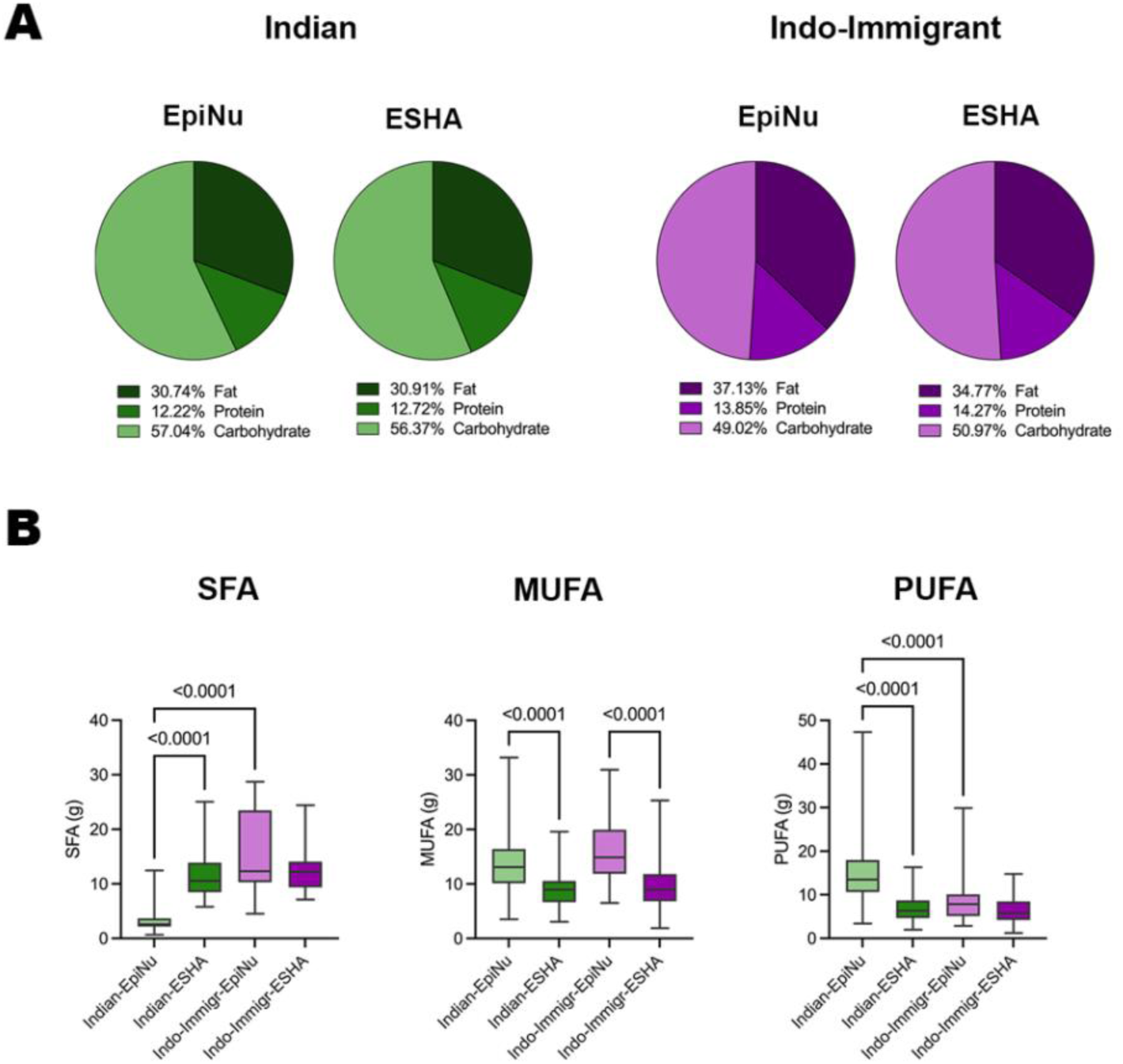
Dietary Fat Types are Significantly Different Between EpiNu vs. ESHA. **(A)** Percentages of fat, protein and carbohydrates were calculated in ESHA and EpiNu for each participant, then mean values were calculated for each cohort and an ordinary one-way Analysis of Variance (ANOVA) was used. **(B)** Significant differences in types of fat consumed by Indians (n = 61) and Indo-Immigrants (n = 32) were calculated between EpiNu vs. ESHA. Dietary fats displayed in grams per 1000 Calories (kcal). Kruskal-Wallis tests were conducted to determine differences between groups. The middle band inside of the boxes is the median, the bottom and top boxes are the first and third quartiles, and the whiskers are the min to max values.

## DISCUSSION

### Westernization is Linked to Both Taxonomic and Functional Transitions in the Gut Microbiomes of Indian Migrants

Earlier studies identified that the Indian gut microbiota is distinct, being enriched in *Prevotella* spp*., Dialister* spp., and *Megasphaera* spp.,^27–33^ unlike westernized groups dominated by *Bacteroides* spp.^34^ and *Phocaeicola vulgatus*^35^. The taxonomic heterogeneity within Indians also highlights how generalizing the “Indian microbiome” may inaccurately represent this population, given the cultural variations that exist within the country. Contrary to associations between low alpha diversity and disease in western populations, healthy Indians showed lower alpha diversity than westernized cohorts, suggesting this diversity measure alone may not always be a reliable marker of westernization or health status.^36^ However, the dominance of *Prevotella spp*. in the Indian gut may have masked the detection of rarer species, potentially underestimating richness estimates in Indians.

Taxonomic data revealed Indians and Indo-Immigrants harbour higher abundances of bacteria characteristic of pre-industrialized societies, including some previously characterized as VANISH taxa. In contrast, Caucasian cohorts possess microbes typical of industrialized populations, including BloSSUM taxa noted in previous studies. However, Indo-Canadians exhibited a transitional microbiota that embodied a blend of both lifestyles, a trend previously detected in infants living in transitional societies (intermediate between non-industrialized and industrialized lifestyles).^37^ A key genus that is distinctive in this transition in immigrants is the loss of *Prevotella* spp.^10,38^ In fact, this contrast in *Prevotella* spp. abundance in non-westernized versus westernized groups has sparked interest both in its relation to lifestyle and disease, previously being linked to IBD,^39,40^ rheumatoid arthritis,^41^ and obesity^42^. Yet, contradictory studies also showed its correlation with reduced risks of cardiovascular disease (CVD) ^43^ diabetes ^44^, and obesity,^45^ illustrating the current contradictory understanding around *Prevotella* spp.-dominance in the gut and its role in diseases.

Globally, Indians have some of the highest abundances of *Prevotella* spp. in their gut,^46^ as reflected in our data, emphasizing the importance of context when investigating the role of microbes in our gut, as host-microbe interactions may divergently evolve depending on the host’s genetics, geography, and lifestyle. In fact, these factors may have already driven evolutionary changes in *Prevotella* spp. strains: isolated *P. copri* strains from Indian/non-western populations showed traits that aid in digestion of plant-based carbohydrates using carbohydrate-active enzymes (CAZymes), whereas *P. copri* strains from westernized populations were enriched with virulence factors and resistance genes.^46^ In our study, Indians had a significantly higher average abundance of *P. copri* Clade A, along with notable abundances of Clades B and C, which subsequently declined in Indo-Immigrants and Indo-Canadians. All *P. copri* clades have previously been documented at higher abundances in non-westernized individuals than westernized,^47^ and our results highlight this effect of westernization on clade abundances using groups of the same ethnicity. These findings demonstrate how specific strains can evolve with its host, cautioning against broad associations between bacterial species and diseases.

Indians, who traditionally consume a high-complex-carbohydrate diet, have been shown to exhibit increased expression of carbohydrate metabolism genes for complex polysaccharides in the gut.^27,31,33^ Our findings reflected this relationship, as we detected enrichment of CAZy families that aid in the digestion of xylan and xyloglucan, with *P. copri* as the top contributing taxa. A similar pattern was in fact detected in another study on immigrants who migrated from Thailand to the US, where a decrease in *Prevotella* spp. abundance corresponded with a loss of CAZymes that break down plant fiber.^10^ The subsequent loss of *P.copri* in our Indo-Immigrant and Indo-Canadian cohorts aligned with their reduction of a high carbohydrate, high fiber diet, which was a trend previously explored among Hadza and Nepali groups.^48^ Plant-derived microbiota-accessible carbohydrates (MACs) are required to maintain abundances of *P. copri*, whereas other species such as *Bacteroides thetaiotaomicron* can utilize both plant and animal-derived MACs, allowing for persistence in the gut.^48^ This relationship suggests that *Prevotella* spp. abundance in the Indian gut is highly linked to their specific dietary patterns. Furthermore, the generational loss of *P. copri* in Indian immigrants prompts questions about gut health implications, meriting further research on strain-specific *P. copri*-host interactions within this demographic.

A previous study found *Dialister succinatiphilus* and *Megasphaera* more abundant in Indo-Immigrants versus Indo-Canadians, who had significantly higher *D. invisus.* Our data align with this trend, with *D. invisus* enriched in Indo-Immigrants, *D. succinatiphilus* in Indo-Canadians, and *Megasphaera* in our Indian cohort. While little is known about these taxa, a relationship between a carbohydrate-rich diet and succinate-utilizing bacteria has been previously discussed,^49^ with *D. succinatiphilus* being a well-known species in this category. Additionally, *Ruminococcus torques,* a known mucin degrader,^50^ previously linked to IBD risk ^51^ and identified as a significant contribution in a Crohn’s Disease microbiome-risk score,^52^ was notably enriched in Indo-Canadians. Deeper investigations should explore the role of specific *R. torques* strains in the gut and its implications in IBD should be explored further.

### India is Currently Industrializing

India’s industrialization is reshaping dietary habits, as reflected in our data showing increased use of sunflower oil over traditional fats like ghee. This trend is likely influenced by globalization^53^ and dietary guidelines recommending a reduction in saturated fat intake and increase in PUFA-rich vegetable oils. Overtime, these vegetable oils became inexpensive and widely accessible in India, contributing to a shift away from traditional fats.^54^ Coinciding with increased PUFA consumption is the rising epidemic of several modern diseases in India such as IBD^55^, type-2 diabetes, and metabolic syndrome.^56^ India is now in an accelerating incidence phase of IBD^1^, doubling the number of patients from 1990 to 2019.^57^ While dietary studies in Indian IBD patients are limited, previous research in other populations has shown that increased intake of linoleic (n-6 PUFA) was associated with an increased risk for UC,^58^ whereas increased intake of α-linolenic acid (n-3 PUFA) was associated with reduced UC-risk.^59^ These patterns highlight the importance of further investigating the potential role of vegetable oil consumption in relation to the rising incidence of IBD in India.

In contrast, the westernized cohorts in our study reported cooking mainly with olive oil and butter, and few reported to cook with vegetable oils rich in omega-6 PUFAs. While these self-reports are subject to limitations, it is worth highlighting that while India has adopted the use of white oils like sunflower oil and canola oil, North America may currently be shifting away from them. Historically, the westernized diet has been characterized by a much higher intake of omega-6 PUFA, leading to omega-6:omega-3 ratios of 20:1, which is far from the estimated ratio of 1:1 of our ancestors.^60^ Excessive omega-6 PUFAs consumption has been associated with increased risks of CVD,^61^ cancer,^62^ and autoimmune diseases.^63^ Furthermore, with large-scale studies and meta-analyses refuting the link between saturated fatty acids and CVD^64,65^ and the growing popularity of the Mediterranean Diet,^66^ a cultural shift in the North America may be occurring, once again preferring olive oil and butter,^67^ which was an observation also recently reported from an Australian cohort.^68^

Another factor of industrialization that is common in India is the overuse of antibiotics. India has one of the highest threats for antimicrobial resistance,^69^ as was reflective in our cohorts with enriched KEGG identities in Indians and Indo-Immigrants that contain signatures for antibiotic resistance genes, a pattern previously documented in the Indian gut.^27,70^ The widespread use of antibiotics in India not only increases the prevalence of antibiotic resistance, but also accelerates microbiome turnover, which was indicative in the pathways enriched in our Indian cohort including peptidoglycan synthesis, microbial growth/metabolism, and DNA synthesis/repair. While common usage of antibiotics is a likely catalyst for cell turnover, the higher risk for infectious disease via water contamination and inadequate sanitation in India may also serve as environmental triggers. Our findings highlight India’s unique transition phase, balancing western influences with traditional lifestyle practices, all of which are shaping their gut microbiome. Future research should delve deeper into how westernization is currently influencing the gut microbiome in Indians, as this will likely pose future health challenges.

### Acculturation of Westernized Diet Drives Functional Shifts in Microbiome

Coinciding with previous findings, we observed dietary acculturation from the traditional high-complex-carbohydrate Indian diet to an increased intake of NOVA Group 4 UPFs,^38,71,72^ which are characterized by industrially made ingredients with non-nutritive additives like emulsifiers, flavouring and sweeteners.^73^ This dietary shift might explain the enrichment of pathways tapping into glycogen storage in Indo-Canadians, suggesting microbes had inadequate nutrient availability from the host diet.^74^ With a 49% rise in Group 4 UPF intake and significant fiber reduction in Indo-Canadians, concerns emerge regarding their gut health and chronic disease risks.

High UPF consumption has been linked to increased risks of CVD, metabolic disorders, micronutrient deficiencies, and anxiety and depression,^73^ which may partly be due to imbalances in the gut microbiome that are a triggered from UPF. While there are several potential culprits involved in the ultra-processing of foods that may have mechanistic interactions with the gut, it is likely an interplay of several factors surrounding the consumption of UPFs. First, despite being affordable and accessible, UPFs are often high in calories and palatability, but low in essential nutrients, leading to overconsumption and displacement of nutritious meals.^75^ Additionally, emulsifiers, used to mix components in UPF, have been extensively studied in murine models, showing their ability to modulate the gut microbiota, reduce mucus thickness and increase intestinal permeability, bacterial translocation and inflammation.^76^ While studies translating these same mechanistic effects of emulsifiers in humans are scarce, a short-term prospective study (*N* = 588) found a positive association between dietary emulsifiers and the inflammatory biomarker glycoprotein acetyls.^77^ Moreover, a randomized double-blind controlled-feeding study (*N* = 16) with the emulsifier carboxymethylcellulose (CMC) found varied responses in the treatment group, with two subjects exhibiting significant alterations in gut composition and microbial encroachment, whereas other participants seemed non-sensitive to CMC.^78,79^ Although constrained by a small sample size, these findings prompt the need to further investigate in large-scale cohorts, as this may in part explain the inconsistent outcomes observed when transitioning from murine to human studies. Overall, these findings underscore that emulsifiers may be one of many components in UPF that influence the gut microbiome, but further human studies are necessary to fully understand the complex role of UPF on the gut microbiome and inflammation. Future research should also consider examining the synergistic effects of multiple food additives to mirror the composition of UPFs typically consumed by humans.

## LIMITATIONS

In addition to inherent limitations of self-reported food diaries (e.g. accuracy of participant reporting), our study also faced limitations in representing cultural diets, primarily due to inadequacies in the existing North American databases for Indian cuisine. This led to the use of the EpiNu Nutritional software, an Indian dietary tool that is tailored for Indian dishes. Another limitation was the shallow depth of shotgun sequencing and taxonomic classification methods used restricted our ability to identify strain-level differences in bacteria like *P. copri*. Uneven sample sizing across groups also presented a limitation; specifically, recruiting Indo-Canadians was considerably more difficult, which may reflect a combination of cultural, social and geographical barriers.

As this project focused on the relative abundance of *Prevotella* spp., quantitative PCR could also help to determine total biomass and absolute abundances of specific taxa present in the gut, as this may also be a key factor influencing disease risk.^80^ Additionally, while metagenomics can provide important insights into the genetic content present within the gut, these data only represent the functional potential of the gut and does not provide information regarding genetic expression and actual function of microbiome communities. Furthermore, BugBase analyzes 16S data against an older version of the Greengenes database, which may influence the accuracy of the predictions, especially in relation to the newer Greengenes2 and MetaPhlAn databases used. Future work should also aim to resolve strain-level analysis as well as parasite communities in the microbiomes of these populations, as this may differ in those living in India versus Canada. Lastly, since this was a cross-sectional study, these data provide associations between westernization and gut microbiome changes, and further investigation is needed to understand the mechanistic interactions of specific factors of westernization on the gut and IBD risk.

## CONCLUSIONS’

Consistent with previous studies, our research reveals that the Indian gut microbiome, enriched in *Prevotella* spp., is adapted to their high carbohydrate, high fiber diet. We discovered that the Indian microbiome exhibits characteristics of higher bacterial cell turnover, pathogenic potential and stress tolerance, which may reflect a chronic alteration in the response to a higher load of pathogen exposure. However, since our subjects were healthy and not displaying GI-related ailments, this may suggest that Indians have a higher resilience against stressors posed on the gut (Figure 8). These findings highlight the importance of studying diverse gut microbiomes, particularly in non-westernized and transitional populations, to better predict and manage diseases like IBD. Our study provides a snapshot on the gut microbiome changes in a population undergoing westernization, revealing a drastic transition within the gut microbiomes of Indo-Immigrants and Indo-Canadians, extending our knowledge of the strong effect of immigration and westernization on the gut microbiome. This transition likely reflects microbial adaptation to the dietary acculturation observed, providing insight into the dynamic nature within our gut. Future investigations should examine the health effects of microbiome changes amidst global westernization, as this transition will likely redefine health outcomes for both migrating populations and those living in rapidly modernizing nations.

**Figure 8.**
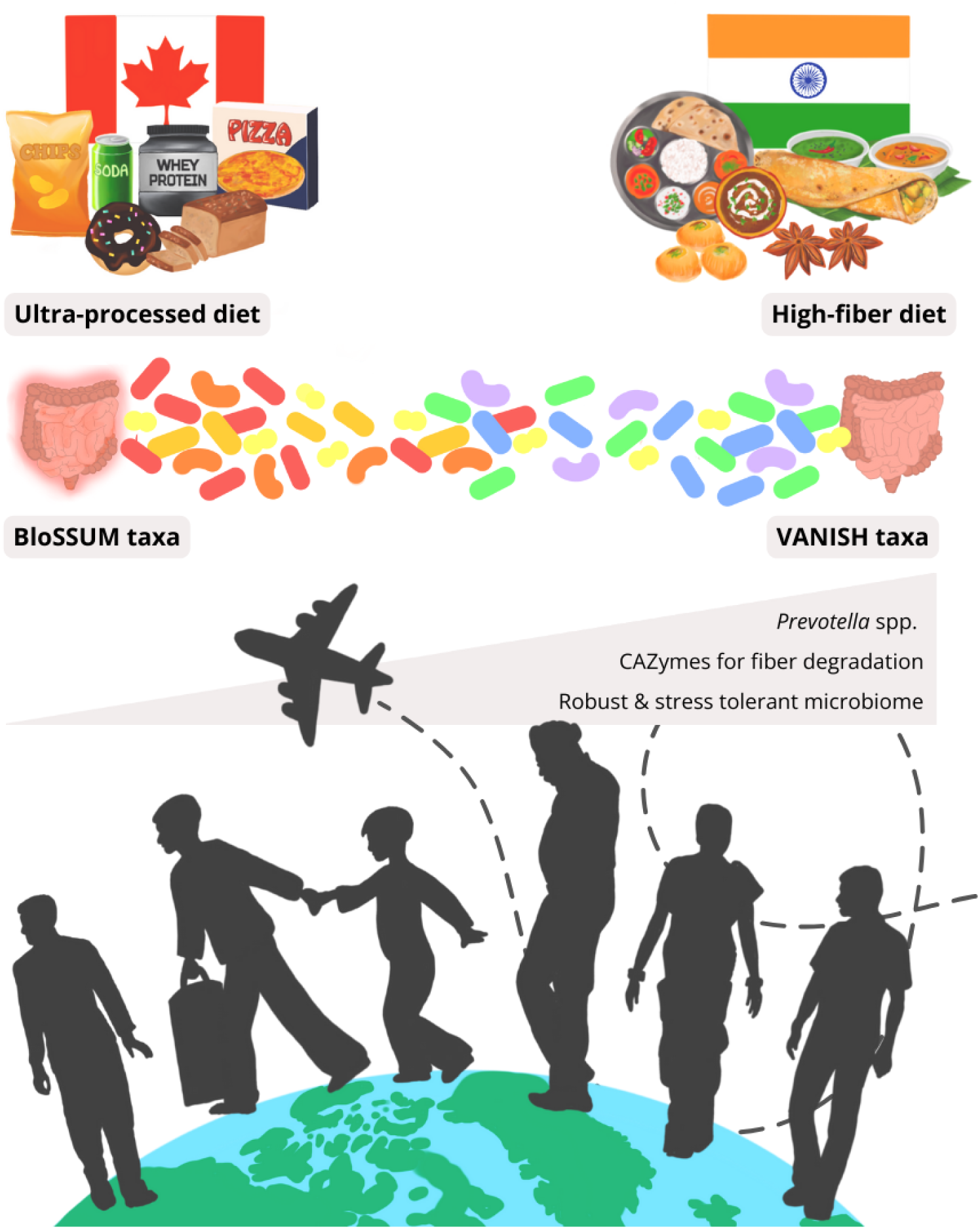
Adoption of Westernized Diet is Associated with a Transition Away from Indian Gut Microbiome. Graphical representation highlighting the main findings from our study. As Indo-Immigrants and Indo-Canadians subsequently increased ultra-processed food intake, their fiber consumption dramatically reduced. This change in dietary pattern was observed alongside a transition away from high *Prevotella* spp., which were found abundant in the Indian gut. Instead, Indo-Canadians adopted BloSSUM (Bloom or Selected in Societies of Urbanization/Modernization) taxa, which are commonly found in the westernized microbiome. High *Prevotella copri* in the Indian gut contributed to enriched carbohydrate-active enzymes (CAZymes) that are specialized to degrade complex carbohydrates common in their diet, which decline in Indian migrant cohorts. Additionally, the Indian microbiome displayed characteristics of a more robust gut, with predictive functions of higher stress tolerance and increased microbial cell turnover.

## Acknowledgements

We thank all the individuals who took part in this study. We thank the entire Ghosh family for support. For dietary analysis, we thank Nadia Anvari. For lab support, we thank Mekenna Smith, Hephzibah Bomide, Carson McComb, Ayva Lewis, Erik Kaila, Andrea Verdugo Meza and Jessica Josephson.

## Funding

This work was supported by the Natural Sciences and Engineering Research Council of Canada (NSERC) and a UBC Killam Research Award to support D.L.G. on her sabbatical in India.

## Competing Interests

DLG is on the Board of Directors for Crohn’s and Colitis Canada and is also the co-founder and CSO of Melius MicroBiomics Inc. where LDD is the Medical Science Writer.

## Patient and Public Involvement

Not applicable.

## Ethics Approval

This study was approved by the University of British Columbia Clinical Research Ethics Board (UBC CREB) (H21-01555). Approval for sample collection in India was approved under IEC No 411/2018 and by UBC CREB (H17-01324). All participants signed a consent form prior to participation. A Southeast Asian (SEA) Microbiome Biobank was created.

## Data Availability Statement

Data supporting this study’s findings are available within the paper and its Supplementary Information. The 16S rRNA gene sequences files and metagenomic sequence files of human fecal samples are publicly available on NCBI under accession number PRJNA1082632.

## Code Availability Statement

The underlying code for this study is available on GitHub and can be accessed via this link: https://github.com/leahdaloisio/India-Microbiome-Project

## Supplementary Material

Please find Supplementary Methods and Supplementary Results attached.

## Notes

https://github.com/leahdaloisio/India-Microbiome-Project

### Competing Interest Statement

DLG is the co-founder and CSO of Melius MicroBiomics where LD is the Medical Science Writer.

### Summary of Updates

Main revisions include: - All figure captions were edited to remove any results statements - Figure 1A-B was updated to show significance values in the boxplots - Figure 2 the taxonomic barplots from the 16S rRNA seq data was removed and added to supplementary. An abundance plot of notable differentially abundant species was added as Figure 2E - Wording was modified (mainly in discussion) to remove superfluous sentences and overconclusions - Github project was made available to include all scripts used for analysis - In supplementary material, the lifestyle survey and immigration stress surveys were included, an abundance plot was added as Figure S4, and Table S1 was added to show average % of unclassified reads per sample in each cohort.

