## Supplementary Material for "The adoption of a westernized gut microbiome in Indian Immigrants and Indo-Canadians is associated with dietary acculturation"

### SUPPLEMENTARY METHODS

#### *Recruitment*

Healthy participants between the ages of 17-55 years were recruited at three sites: (1) Kolkata, Indian (2) Manipal, India (3) Kelowna, Canada. In Canada, participants were recruited from the University of British Columbia Okanagan campus, along with religious temples in Kelowna and Vancouver. Posters and pamphlets were handed out to the community, with recruitment material tailored for both Punjabi and Indian/Bengali communities. Participants were excluded if they were pregnant or had a diagnosis of any chronic inflammatory condition. Samples also were not collected if participants had taken antibiotics or travelled to India less than three months prior.

#### *Lifestyle Analysis*

To determine immigration-related stress from Indo-Immigrants and Euro-Immigrants, a survey was provided that asked questions regarding their experiences since migration to Canada, such as stress involving documentation, financial insecurity, discrimination, etc. Questions were adopted from Sternberg 2016, in which they administered a survey to Mexican immigrants in the United States,[1] with questions removed/modified to be applicable for Canadian immigrants in our study (see below). While it can be assumed that since Euro-Immigrants were already westernized, they would experience less of a change in lifestyle in Canada compared to Indo-Immigrants. We wanted to confirm this through the completion of a survey in which subjects were provided a list of statements of lifestyle changes towards westernization, and they were instructed to score how relevant these statements were to their experience (see below).

a place of mind
THE UNIVERSITY OF BRITISH COLUMBIA

Participant ID: ____________________

Date: ___________________________

IMMIGRATION STRESS

Please check the answer that best describes how relevant each statement applies to your life **since immigrating to Canada.**

1 = LOW STRESS    2 = MODERATE STRESS    3 = HIGH STRESS

| Q | Statement | 1 | 2 | 3 |
|---|---|---|---|---|
| 1 | How much stress/worry do you experience because documentation problems (i.e. Canadian citizenship, PR cards) keep you from getting the things that you need for you and your family? | | | |
| 2 | How much stress/worry do you experience because documentation (i.e. Canadian citizenship, PR cards) problems make it difficult for you to visit your country? | | | |
| 3 | How much stress/worry do you experience because you feel you cannot compete with Canadians in your workplace? | | | |
| 4 | How much stress/worry do you experience because of financial insecurity and/or economic stress? | | | |
| 5 | How much stress/worry do you experience because you miss your family and friends back in your home country? | | | |
| 6 | How much stress/worry do you experience because you feel emotional or sentimental when thinking about your life back in your home country? | | | |
| 7 | How much stress/worry do you experience because you feel it is hard to face new situations and circumstances here in Canada (e.g. renting an apartment)? | | | |
| 8 | How much stress/worry do you experience because you feel that cultural differences in Canada are causing conflicts within your family? | | | |
| 9 | How much stress/worry do you experience because you feel people discriminate against you and you are treated as a second-class citizen? | | | |
| 10 | How much stress/worry do you experience because you feel that this is not your country although you live here? | | | |
| **Do you have any additional comments you’d like to make regarding the stress you experience as an immigrant in Canada?** | | | | |

Questions from: Sternberg, R. M., Nápoles, A. M., Gregorich, S., Paul, S., Lee, K. A., & Stewart, A. L. (2016). Development of the Stress of Immigration Survey (SOIS): A field test among Mexican immigrant women. *Family & community health*, *39*(1), 40.

a place of mind
THE UNIVERSITY OF BRITISH COLUMBIA

Participant ID: ____________________

Date: ___________________________

Think about how you lived in your homeland versus living in Canada.
Answer each question in the context of **how your life has changed since living in Canada VERSUS living in India/Punjab/Bangladesh.**

Please check the answer that best describes how relevant each statement applies to your life:

| Q | Statement | Strongly Agree | Agree | Neutral | Disagree | Strongly Disagree |
|---|---|---|---|---|---|---|
| 1 | My consumption of fast food has increased since moving to Canada<br>E.g. McDonalds, KFC, Domino's Pizza, A&W | | | | | |
| 2 | My consumption of processed foods has increased since moving to Canada<br>E.g. microwave meals, chips, instant noodles, frozen pizza, sugary cereals | | | | | |
| 3 | My consumption of sugary drinks has increased since moving to Canada.<br>E.g. Coke, Sprite, Gatorade, RedBull | | | | | |
| 4 | The proportion of plant-based foods in my diet has decreased since moving to Canada.<br>E.g. vegetables, fruits & legumes, grains | | | | | |
| 5 | My consumption of psychotropic medications has increased since moving to Canada.<br>E.g. antidepressants, anti-anxiety, mood stabilizers, antipsychotic, stimulants | | | | | |
| 6 | I have become more sedentary since moving to Canada.<br>E.g. spend more time sitting, laying down, less physically active overall | | | | | |
| 7 | My sanitation/hygiene practices have increased since moving to Canada<br>E.g. Better water quality, hygiene practices | | | | | |
| 8 | My level of emotional distress has increased since moving to Canada<br>E.g. anxiety, depression, overwhelmed | | | | | |

#### *Stool Collection & DNA Extraction*

Participants were given at-home stool collection kits consisting of a large plastic container (Medline, 320-DYND36500), a stool collection hat (Medline, DYND36600), gloves, a facemask, and a stool collection guide. Subjects stored their sample in their freezer for no longer than 3 days. Samples were transported on dry ice to the lab, then stored in a -80°C freezer until homogenization. In a biosafety cabinet, stool was homogenized in liquid nitrogen, then stored in -80°C. DNA was extracted from stool samples using the QIAamp PowerFecal Pro DNA Kit (Qiagen, Cat. No. 51804) following the manufacturer’s instructions. All samples except those collected in India had an additional wash step (C5) to improve DNA purity. DNA samples were sent to Gut4Health Microbiome Core Facility (BC Children’s Hospital Research Institute, Vancouver, British Columbia) for 16S sequencing on the Illumina MiSeq platform (V4-V4 region amplified with 515f and 806r primers) (~75,800 reads per sample). For shotgun sequencing, DNA concentration was normalized to 30uL in nuclease free water, then sent to the Center for Health Genomics and Informatics (University of Calgary, British Columbia) for shotgun sequencing on the Illumina NovaSeq platform (~9.9M reads per sample).

#### *16S Microbiome Analysis*

Paired-end demultiplexed reads were imported into QIIME 2 (Version 2022.2) [2]. Quality control was completed with DADA2, which included filtering, chimera removal, dereplication, denoising and merging paired-end reads [3]. For taxonomic classification, the q2-feature-classifer [4] was trained using the GreenGenes2 database (10.28.22)[5]. Amplicon sequence variants (ASVs) were filtered for unclassified ASVs (i.e., identified only to phyla level), sequence counts below 1000, and mitochondrial/chloroplast DNA. Using q2-alignment, ASVs were aligned with mafft [6] to construct a phylogeny with fastree via q2-phylogeny [7]. Alpha diversity was calculated with Shannon [8] and Pielou’s Evenness, both diversity metrics were also calculated with rarefied data at a sampling depth of 12053, but no major differences were observed. [9]. Beta diversity metrics were calculated using rarefied data with Bray Curtis [10] and Weighted UniFrac [11]. To determine if there were differentially abundant bacteria across cohorts, the Linear Discriminate Analysis (LDA) Effect Size (LEfSe) algorithm was used via the Huttenhower Lab Galaxy Hub, [12] using a one-against-all strategy with a threshold of 3.5 on the logarithmic LDA score and an alpha of 0.05 for Kruskal-Wallis test among classes. BugBase was used to predict microbiome phenotypes such as Gram-negative, Gram-positive, potentially pathogenic and stress-tolerant bacteria.[13-17] To understand associations between the distinctions in beta diversity from the taxonomic data with the dietary patterns and baseline characteristics that were significantly different, a distance-based redundancy analysis (dbDRA) was executed in R statistical software (4.2.2),[18] using the Weighted UniFrac plot generated from 16S amplicon data in QIIME 2.

#### *Shotgun Taxonomic Analysis*

Paired-end demultiplexed raw sequence reads were first assessed for quality using FastQC and all samples were compiled into a MutliQC report to determine parameters for trimming. The input sequence data had an average of 9.9 million reads per sample, with an average length of 127 base pairs. Quality control was performed using KneadData (Version 0.10), with reads trimmed based on the minimum average quality threshold of 25 with a search-window size of 4 bases (SLIDINGWINDOW:4:25). To additionally improve quality, a fixed number of 10 base pairs were trimmed from the ends of reads, and any sequences exceeding a length of 120 base pairs were trimmed down to this maximum length. Reads shorter than the minimum threshold of 60 base pairs were removed. Additionally, host contamination, primers and sequencing adapters were also removed using KneadData. Human host sequences were removed by aligning reads against a host genome database and eliminating perfectly mapped reads.

Taxonomic profiling was conducted using MetaPhlAn4, then each sample was normalized to relative abundances. The normalized relative abundance output table was then imported into R statistical software for downstream analysis including alpha diversity metrics Shannon [8] and Pielou's Evenness [9], as well as beta diversity metrics Bray Curtis [10] and Weighted UniFrac [11]. Differentially abundant bacteria were determined using LEfSE with a 3.5 threshold and alpha 0.05 [12]. The normalized abundance output table was imported into MicrobiomeAnalyst 2.0 to generate a heatmap of highest abundance taxa across cohorts [19]. For the heatmap, a low count filter was applied to remove features less than 4 in fewer than 20% of samples, and a low variance filter was set to remove the lowest 10% of features with minimal variability, as determined by an inter-quantile range threshold.

***Nutritional Analysis***

Participants filled out a food log for 3 consecutive days prior to their stool collection. Food logs were then entered in the ESHA Food Processor® software, which generated a table for each participant with their nutrient intake for each day. Several data cleaning steps were untaken to ensure food intake was inputted into ESHA to accurately reflect the participant's food log (i.e. calories, proportions, recipes, etc.). One researcher had conducted 4 rounds of data cleaning and three additional researchers reviewed ESHA data for outliers such as abnormally high caloric or nutrient intake. Each nutrient was averaged over 3 days, and results were reported both as absolute values (males and females separate) and as values normalized to 1000 calories. To account for potential biases in using a North American nutritional software, we inputted food logs into the Madras Diabetes Research Foundation (MDRF) EpiNu® Nutritional software, which was designed to represent the Indian diet. If subjects did not specify the type of cooking oils, the cooking oil type was adjusted to reflect the dominant cooking oil used in the subject's region.

To calculate the percentage of daily caloric intake from ultra-processed foods (UPFs), dietary data for each individual was examined in ESHA to flag any products that were classified as UPF according to the NOVA classification. Calories from UPFs were added up for each participant, averaged over 3 days, then divided by the average caloric intake of each participant x 100.

To determine participants who were vegetarian or pescetarian, participants were asked in the demographic questionnaire if they had any dietary restrictions. In addition to the subject's reported diet, the food logs themselves were scanned for validation. Pescatarians were counted if their food logs reported to only be eating fish, but not meat. Whereas vegetarians were counted if their food records did not contain any meat, fish, but did include eggs (ovo-vegetarian) and/or dairy (lacto-vegetarian).

***Power Analysis***

To estimate the required sample size per group, beta diversity distance matrix scores were used.[20] Due to the nonparametric nature of our data, median and IQR values were used instead of mean and standard deviation. The required sample size was determined using the following formula:

$$n = 2\,\frac{\left(\mathrm{Z}\frac{\alpha}{2} + \mathrm{Z}_{1-\beta}\right)^2}{\Delta^2} \qquad \Delta = \frac{M_1 - M_2}{IQR}$$

Median Bray Curtis distances of 0.8273044 and 0.608826 for the Indian and Indo-Immigrant cohorts were used, respectively, and the larger IQR (0.1767818) was chosen. Therefore, with a

5% alpha error and a 20% beta error, the total estimated sample size required per group was 10 participants.

### SUPPLEMENTARY RESULTS

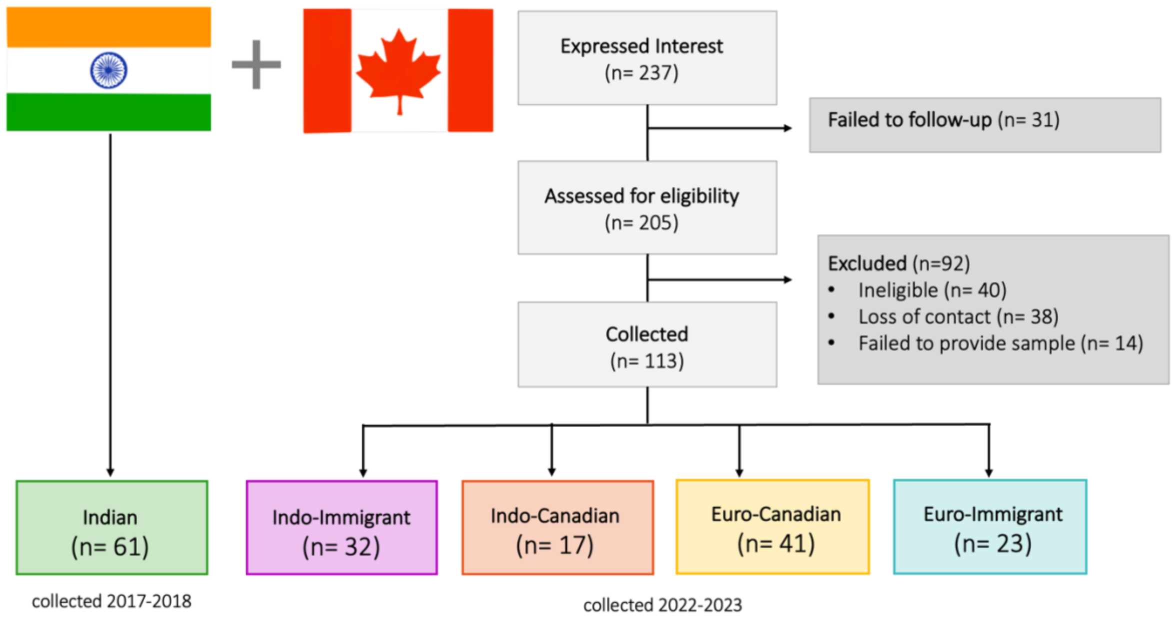


**Figure S1. Recruitment flow chart**

Participants were recruited from Kolkata and Manipal (India) and Kelowna (Canada). Total samples collected from each cohort were Indian (*n* = 61), Indo-Immigrant (*n* = 32), Indo-Canadian (*n* = 17), Euro-Canadian (*n* = 41), Euro-Immigrant (*n* = 23). Samples were first collected in 2017-2018 in India, then due to the pandemic, recruitment did not begin in Canada until 2022.

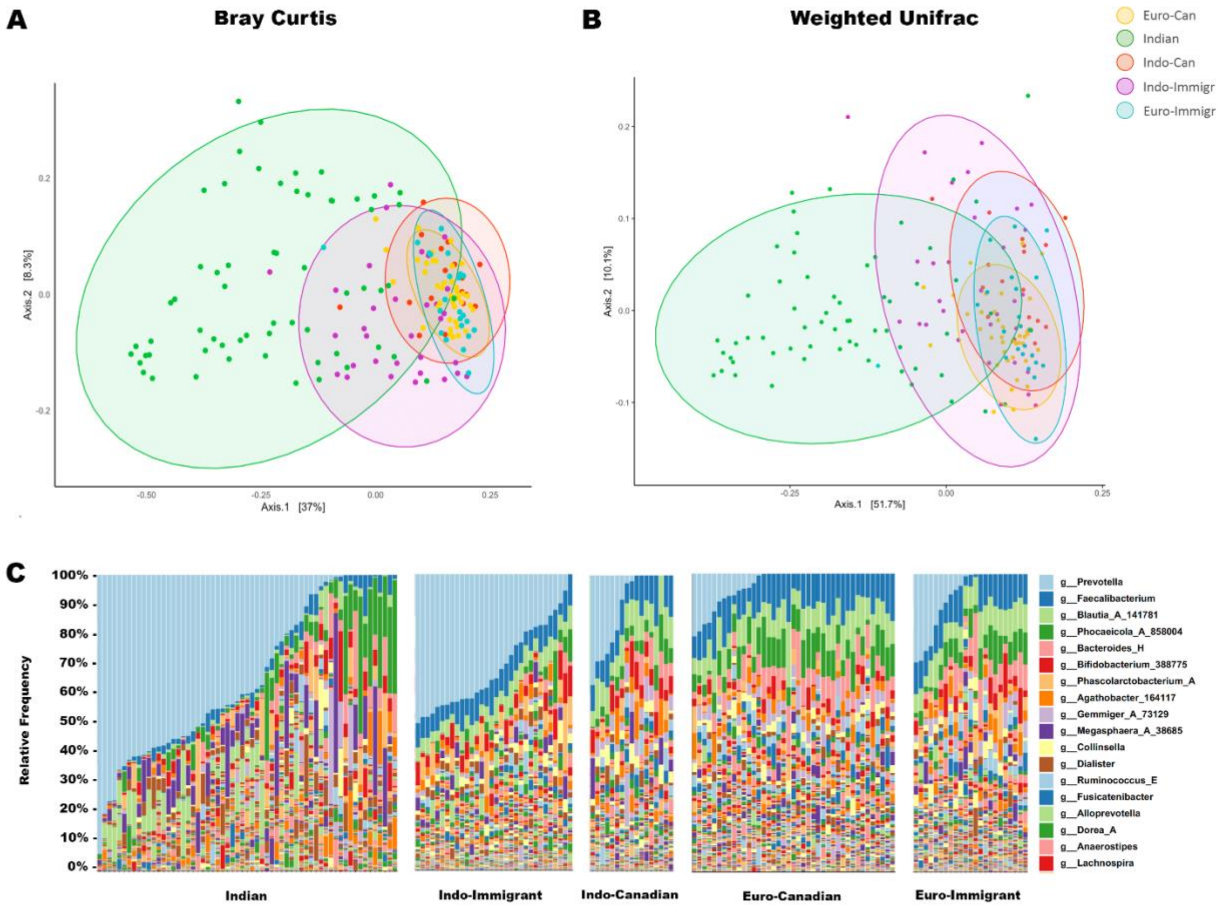


**Figure S2. Beta diversity measures of bacterial communities in study cohorts from 16S amplicon data**

Using 16S amplicon sequences, demultiplexed forward and reverse reads were truncated at 251 bases. Beta diversity was explored with Bray Curtis dissimilarity and Weighted UniFrac, using Pairwise Permutational Multivariate Analysis of Variance (PERMANOVA) to test differences between groups. **(A)** Bray Curtis principal coordinate analysis (PCoA) plot shows 28.6% of variation was captured on the first two axes. Pairwise comparisons with Bonferroni correction revealed significant differences between most groups, with distinct clustering of Indians and Indo-Immigrants from westernized cohorts. Indians show significant distinctions in their gut microbiota when compared to Indo-Immigrants (pseudo-F = 6.585, $p_{BONF}$ = 0.001), Indo-Canadians (pseudo-F = 8.203, $p_{BONF}$ = 0.001), and Euro-Canadian controls (pseudo-F = 21.60, $p_{BONF}$ = 0.001). Indo-Immigrants were also significantly distinct from Indo-Canadians (pseudo-F = 6.094, $p_{BONF}$ = 0.001). Indo-Canadians were significantly distinct from Euro-Canadians (pseudo-F = 2.918, $p_{BONF}$ = 0.001). **(B)** Weighted UniFrac PCoA plot shows 58.3% of the variation was captured on the first two axes, with significant differences also detected. Indians were significantly distinct from Indo-Immigrants (pseudo-F = 7.707, $p_{BONF}$ = 0.001), Indo-Canadians (pseudo-F = 15.95, $p_{BONF}$ = 0.001), and Euro-Canadians (pseudo-F = 49.18, $p_{BONF}$ = 0.001). Indo-Canadians were also distinct from Indo-Immigrants (pseudo-F = 12.86, $p_{BONF}$ = 0.001) and Euro-Canadians (pseudo-F = 6.748, $p_{BONF}$ = 0.001). **(C)** Taxonomic stacked bar plots depicting relative abundances of top 18 most abundant bacteria across all samples from 16S rRNA sequence data (N = 174; Indian (n = 61), Indo-Immigr (n = 32), Indo-Can (n = 23), Euro-Can (n = 41), Euro-Immigr (n = 23).

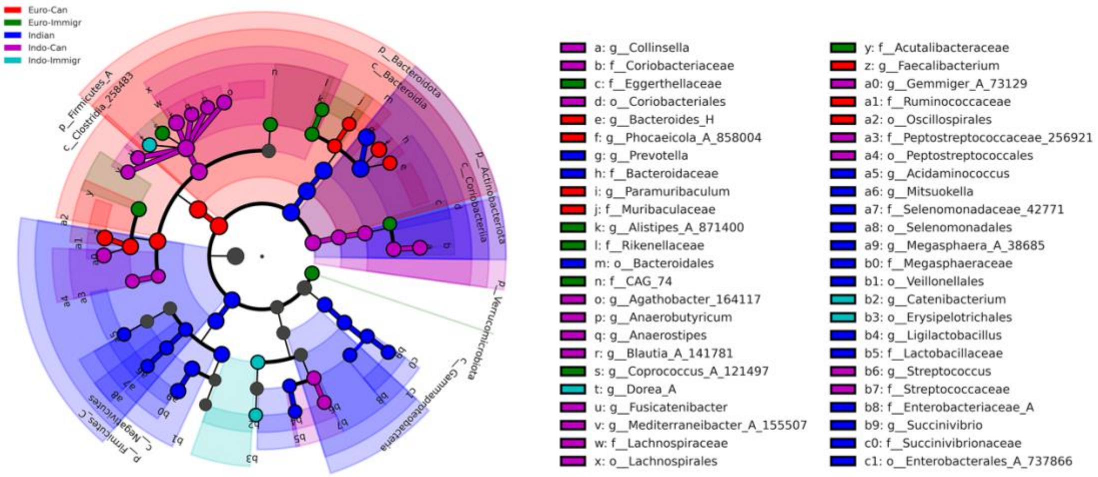


**Figure S3. LEfSe cladogram results from 16S amplicon data**

LEfSe (Linear discriminate analysis Effect Size) cladogram results, depicting differentially abundant bacteria across cohorts, with genus set as lowest taxonomic rank. Indian gut displayed a dominance of Bacteroidota, Gammaproteobacteria and Firmicutes_C, Euro-Canadians showed high abundance of Firmicutes_A. Indo-Immigrants had high abundances of Catenibacterium, Erysipelotrichales and Dorea_A. Indo-Canadians predominated with Actinobacteria.

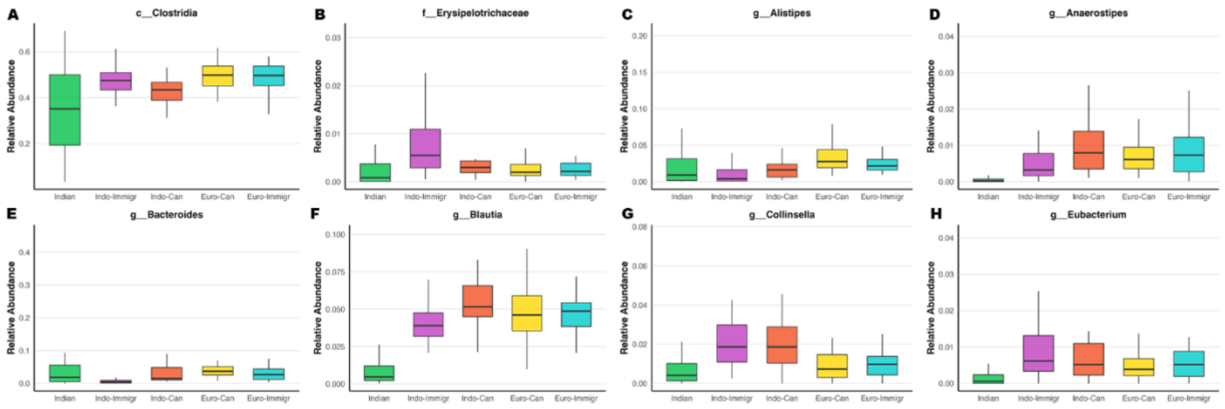


**Figure S4. Average relative abundance plots of differentially abundant taxa identified by LEFSE**

Boxplots depicting average relative abundance of taxa previously identified common in pre-industrialized **(B,G,H)** and industrialized microbiomes **(A, C, D, E, F).**

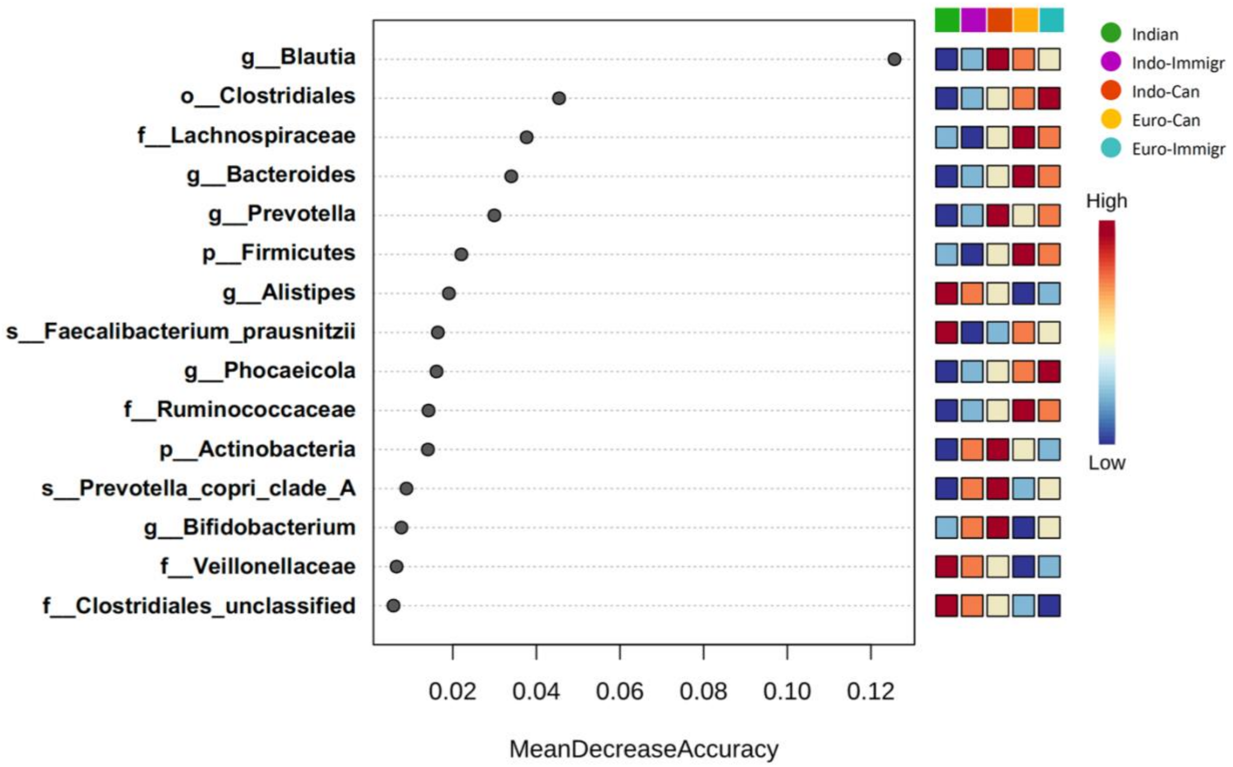


**Figure S5. Random forest results from shotgun data**

Random Forest results generated in MicrobiomeAnalyst 2.0 using MetaPhlAn4 relative abundance table output. Features are ranked by their contributions to classification accuracy (Mean Decrease Accuracy), indicating the extent to which the presence/absence of each bacteria influences the overall accuracy of classifying samples into their respective cohorts.

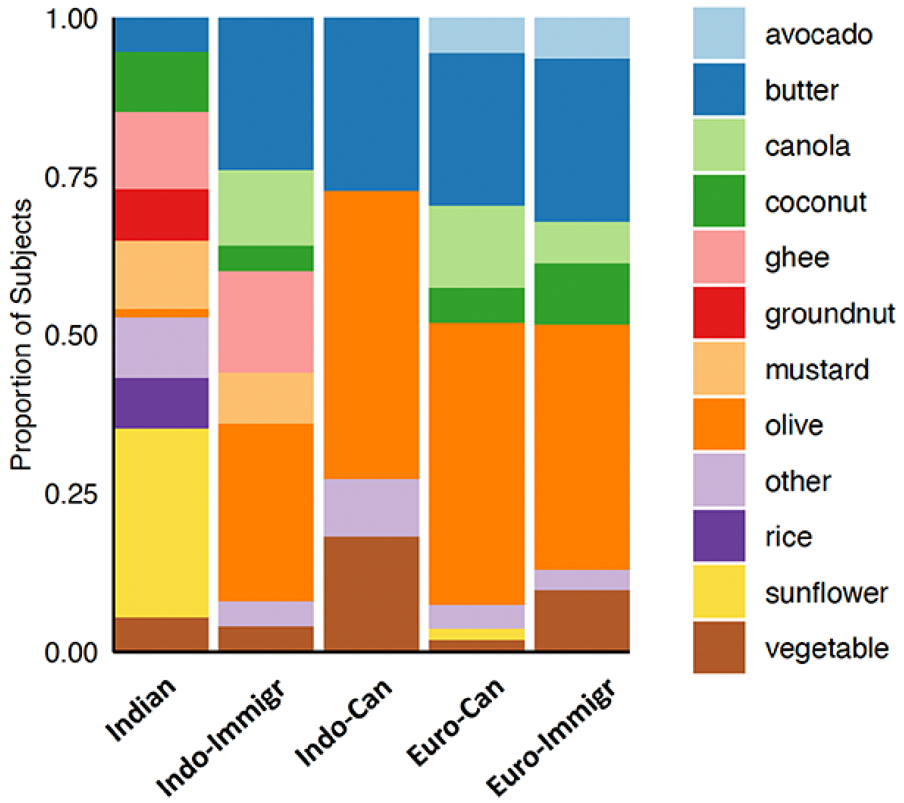


**Figure S6. Decreased variability in cooking oils are used by Indian migrants**

Food logs were reviewed for participants who reported their use of cooking oils/fats. A total of 70% (*n* = 43/61) of Indians, 56% (*n* = 18/32) of Indo-Immigrants, 53% (*n* = 9/17) Indo-Canadians, 85% (*n* = 35/41) Euro-Canadians, and 78% (*n* = 18/23) of Euro-Immigrants reported the types of fats they used when cooking. This plot displays the number of subjects in each group who reported their use of the listed cooking oils/fats. Oils that were reported less than 5 times across all cohorts were grouped into the “other” category. In Indians, the most dominant cooking fats reported were sunflower oil and ghee. Compared to Indians, both Indian migrant cohorts reported much less variety in cooking oils. Indo-Immigrants, Indo-Canadians , and both westernized control groups predominately used olive oil and butter.

**Table S1. Average percentage of unclassified reads per sample per cohort**

| Group | 16S rRNA<br>% unclassified | Shotgun<br>% unclassified |
|---|---|---|
| **Indian** | 1.85% | 6.13% |
| **Indo-Immigr** | 0.12% | 4.47% |
| **Indo-Can** | 1.18% | 3.47% |
| **Euro-Can** | 0.15% | 4.80% |
| **Euro-Immigr** | 0.13% | 3.77% |

**Table S2. LEfSe LDA results from 16S amplicon data**

| Taxon | Log Score | Class | LDA | P value |
|---|---|---|---|---|
| *Succinivibrio sp.* (000431835) | 4.212 | Indian | 3.919 | 9.73E-05 |
| *Prevotella hominis* | 4.400 | Indian | 4.087 | 1.34E-07 |
| *Prevotella stercorea* | 4.236 | Indian | 3.883 | 1.08E-10 |
| *Prevotella copri* | 5.426 | Indian | 5.103 | 2.12E-17 |
| *Megasphaera* A (38685) | 4.820 | Indian | 4.508 | 5.08E-15 |
| *Dialister hominis* | 4.141 | Indian | 3.832 | 5.98E-06 |
| *Mitsuokella multacida* | 3.832 | Indian | 3.535 | 8.44E-14 |
| *Ligilactobacillus ruminis* | 3.872 | Indian | 3.600 | 2.37E-18 |
| *Acidaminococcus* | 3.937 | Indian | 3.584 | 2.83E-08 |
| Catenibacterium sp. (000437715) | 3.880 | Indo-Immigr | 3.555 | 2.15E-10 |
| *Dorea* A *longicatena* | 4.144 | Indo-Immigr | 3.720 | 6.21E-14 |
| *Dialister succinatiphilus* | 4.022 | Indo-Immigr | 3.754 | 0.000355 |
| *Anaerostipes hadrus* | 4.310 | Indo-Can | 3.964 | 8.58E-20 |
| *Anaerobutyricum* | 4.093 | Indo-Can | 3.737 | 2.33E-20 |
| *Streptococcus* | 4.087 | Indo-Can | 3.681 | 9.80E-08 |
| *Blautia* A (141781) | 4.759 | Indo-Can | 4.403 | 2.25E-24 |
| *Fusicatenibacter saccharivorans* | 4.371 | Indo-Can | 4.028 | 2.86E-17 |
| *Mediterraneibacter* A (155507) *faecis* | 4.048 | Indo-Can | 3.695 | 2.93E-11 |
| *Gemmiger* A (73129) | 4.428 | Indo-Can | 4.039 | 1.35E-08 |
| *Blautia* A (141781) *massiliensis* | 4.413 | Indo-Can | 4.054 | 1.06E-14 |
| *Collinsella* | 4.539 | Indo-Can | 4.080 | 4.80E-09 |
| *Agathobacter rectalis* | 4.607 | Indo-Can | 4.162 | 3.44E-05 |
| *Mediterraneibacter* A (155507) | 4.240 | Indo-Can | 3.902 | 1.48E-17 |
| Peptostreptococcaceae (256921) | 4.067 | Indo-Can | 3.509 | 5.83E-09 |
| *Faecalibacterium prausnitzii* C (71358) | 4.755 | Euro-Can | 4.285 | 6.92E-13 |
| *Blautia* A (141781) *faecis* | 4.165 | Euro-Can | 3.826 | 2.80E-21 |
| *Paramuribaculum* sp. (900551515) | 3.962 | Euro-Can | 3.579 | 4.55E-08 |
| *Bacteroides* H | 4.268 | Euro-Can | 3.893 | 2.75E-09 |
| *Phocaeicola* A (858004) *vulgatus* | 4.728 | Euro-Can | 4.359 | 2.12E-08 |
| *Gemmiger* A (73129) *qucibialis* | 4.207 | Euro-Can | 3.800 | 3.94E-10 |
| *Faecalibacterium* | 5.026 | Euro-Can | 4.607 | 3.71E-20 |
| *Alistipes* A (871400) | 4.077 | Euro-Immigr | 3.696 | 1.68E-13 |
| *Coprococcus* A (121497) | 4.257 | Euro-Immigr | 3.869 | 0.010928 |

**Table S3. Differential microbial metabolic pathways across cohorts**

| Pathway | Log Score | Class | LDA | P value |
|---|---|---|---|---|
| Superpathway of L-Aspartate and L-Asparagine Biosynthesis | 3.758 | Indian | 3.067 | 2.09E-13 |
| Glycolysis I (from Glucose-6-Phosphate) | 3.739 | Indian | 3.091 | 3.70E-11 |
| Peptidoglycan Biosynthesis I (Meso-Diaminopimelate Containing) | 4.134 | Indian | 3.138 | 0.000606 |
| 3-Deoxy-D-Manno-Octulosonate Biosynthesis | 3.647 | Indian | 3.179 | 3.37E-17 |
| Folate Transformations II (Plants) | 4.057 | Indian | 3.001 | 0.000615 |
| Inosine 5-Phosphate Degradation | 4.059 | Indian | 3.052 | 1.49E-07 |
| Cis-Vaccenate Biosynthesis | 3.690 | Indian | 3.022 | 1.73E-09 |
| Superpathway of Guanosine Nucleotides De Novo Biosynthesis II | 3.706 | Indian | 3.076 | 5.52E-10 |
| Superpathway of Adenosine Nucleotides De Novo Biosynthesis II | 3.778 | Indian | 3.091 | 5.73E-10 |
| Hydroxymethyl-Dihydropterin Diphosphate Biosynthesis I | 3.693 | Indian | 3.171 | 3.78E-12 |
| Peptidoglycan Biosynthesis III (Mycobacteria) | 4.127 | Indian | 3.236 | 4.29E-06 |
| UDP-N-Acetylmuramoyl-Pentapeptide Biosynthesis II (Lysine Containing) | 4.141 | Indian | 3.119 | 0.001619 |
| UDP-N-Acetylmuramoyl-Pentapeptide Biosynthesis I (Meso-Diaminopimelate | 4.138 | Indian | 3.149 | 0.000520 |
| Queuosine Biosynthesis I (De Novo) | 4.096 | Indian | 3.276 | 7.39E-07 |
| Pre Q0 Biosynthesis | 3.821 | Indian | 3.183 | 1.54E-11 |
| Pyrimidine Deoxyribonucleosides Salvage | 3.968 | Indian | 3.223 | 2.97E-06 |
| Superpathway of Pyrimidine Nucleobases Salvage | 3.655 | Indian | 3.011 | 4.06E-08 |
| Guanosine Ribonucleotides De Novo Biosynthesis | 4.185 | Indian | 3.311 | 6.22E-06 |
| Superpathway of Guanosine Nucleotides De Novo Biosynthesis I | 3.765 | Indian | 3.118 | 2.33E-10 |
| Superpathway of Adenosine Nucleotides De Novo Biosynthesis I | 3.868 | Indian | 3.117 | 1.42E-10 |
| UDP-N-Acetylmuramoyl-Pentapeptide Biosynthesis III (Meso-Diaminopimelate) | 4.111 | Indian | 3.216 | 5.60E-06 |
| Peptidoglycan Maturation (Meso-Diaminopimelate Containing) | 3.957 | Indian | 3.148 | 1.04E-07 |
| L-Valine Biosynthesis | 4.135 | Indian | 3.024 | 0.001861 |
| L-Arginine Biosynthesis I (via L-Ornithine) | 4.010 | Indo-Immigr | 3.229 | 5.15E-17 |
| L-Ornithine Biosynthesis I | 4.015 | Indo-Immigr | 3.361 | 3.48E-22 |
| Thiamine Phosphate Formation from Pyrithiamine and Oxythiamine (Yeast) | 3.950 | Indo-Immigr | 3.037 | 1.73E-08 |
| Superpathway of Thiamine Diphosphate Biosynthesis III (Eukaryotes) | 3.691 | Indo-Immigr | 3.065 | 8.88E-15 |
| Superpathway of Adenosylcobalamin Salvage from Cobinamide I | 3.831 | Indo-Can | 3.360 | 3.32E-22 |
| Pentose Phosphate Pathway (Non-Oxidative Branch I) | 3.887 | Indo-Can | 3.098 | 5.27E-10 |
| Pyruvate Fermentation to Acetate and (S)-Lactate I | 3.676 | Indo-Can | 3.018 | 3.48E-16 |
| Glycogen Degradation II | 4.067 | Indo-Can | 3.369 | 9.27E-20 |
| D-Galactose Degradation I (Leloir Pathway) | 3.883 | Indo-Can | 3.126 | 2.96E-18 |
| Molybdopterin Biosynthesis | 3.867 | Indo-Can | 3.359 | 1.32E-22 |
| Pentose Phosphate Pathway (Non-Oxidative Branch II) | 3.922 | Indo-Can | 3.039 | 1.07E-11 |
| Diacylglycerol Biosynthesis I | 3.941 | Euro-Can | 3.166 | 2.16E-15 |

| | | | | |
|---|---|---|---|---|
| Adenine and Adenosine Salvage III | 4.166 | Euro-Can | 3.070 | 0.000548 |
| Purine Ribonucleosides Degradation | 4.074 | Euro-Can | 3.401 | 5.14E-18 |
| Diacylglycerol Biosynthesis II | 3.941 | Euro-Can | 3.166 | 2.16E-15 |
| Fatty Acid Biosynthesis Initiation (Mitochondria) | 4.045 | Euro-Can | 3.403 | 1.07E-21 |
| L-Arginine Biosynthesis II (Acetyl Cycle) | 4.032 | Euro-Immigr | 3.344 | 2.48E-20 |
| Glycogen Biosynthesis I (from ADP-D-Glucose) | 4.117 | Euro-Immigr | 3.468 | 1.52E-21 |
| Aminoimidazole Ribonucleotide Biosynthesis I | 4.040 | Euro-Immigr | 3.206 | 9.73E-19 |
| Isoprene Biosynthesis I | 3.927 | Euro-Immigr | 3.310 | 2.74E-18 |
| Sucrose Biosynthesis II | 4.183 | Euro-Immigr | 3.555 | 4.44E-21 |
| Arginine Biosynthesis IV (Archaebacteria) | 3.769 | Euro-Immigr | 3.101 | 3.81E-09 |
| Flavin Biosynthesis I (Bacteria and Plants) | 3.898 | Euro-Immigr | 3.162 | 2.23E-17 |

**Table S4. Differential abundances of CAZyme genes**

| CAZyme | Log Score | Class | LDA | P value |
|---|---|---|---|---|
| GH10 | 4.243 | Indian | 3.921 | 1.37E-18 |
| GH106 | 4.100 | Indian | 3.596 | 3.24E-05 |
| GH110 | 3.968 | Indian | 3.628 | 1.18E-09 |
| GH28 | 4.272 | Indian | 3.724 | 0.000284 |
| GH43 | 5.017 | Indian | 4.417 | 1.57E-11 |
| GH51 | 4.586 | Indian | 4.047 | 2.57E-11 |
| GH53 | 3.882 | Indian | 3.414 | 0.000149 |
| GT19 | 3.662 | Indian | 3.220 | 0.003873 |
| GT51 | 4.357 | Indian | 3.866 | 1.13E-10 |
| GT83 | 3.530 | Indian | 3.204 | 4.40E-18 |
| PL1 | 4.235 | Indian | 3.858 | 3.30E-09 |
| GH1 | 4.351 | Indo-Immigr | 3.922 | 8.19E-14 |
| CBM48 | 4.640 | Indo-Can | 4.094 | 1.97E-13 |
| GH113 | 3.549 | Indo-Can | 3.223 | 3.11E-07 |
| GH13 | 4.995 | Indo-Can | 4.381 | 7.49E-12 |
| GH31 | 4.305 | Indo-Can | 3.801 | 3.41E-11 |
| GH4 | 3.882 | Indo-Can | 3.365 | 2.79E-07 |

| | | | | |
|---|---|---|---|---|
| GH42 | 3.811 | Indo-Can | 3.300 | 1.36E-05 |
| GH77 | 4.464 | Indo-Can | 3.944 | 3.66E-14 |
| GH78 | 4.074 | Indo-Can | 3.474 | 9.43E-08 |
| GT28 | 4.477 | Indo-Can | 3.767 | 2.37E-06 |
| GT35 | 4.497 | Indo-Can | 4.019 | 1.28E-15 |
| GH109 | 4.300 | Euro-Can | 3.897 | 1.03E-10 |
| GH23 | 4.425 | Euro-Can | 3.706 | 1.49E-05 |
| GH88 | 3.898 | Euro-Can | 3.438 | 1.96E-07 |
| GH92 | 4.248 | Euro-Can | 3.827 | 1.03E-09 |
| GT2 | 5.072 | Euro-Can | 4.359 | 8.34E-11 |
| GT30 | 3.587 | Euro-Can | 3.065 | 3.26E-06 |
| GT4 | 4.571 | Euro-Can | 3.705 | 0.012273 |
| GT5 | 4.397 | Euro-Can | 3.858 | 2.33E-13 |
| PL10 | 3.837 | Euro-Can | 3.421 | 1.19E-08 |
| PL12 | 3.557 | Euro-Can | 3.169 | 2.91E-08 |
| PL8 | 3.580 | Euro-Can | 3.148 | 3.04E-06 |
| CBM13 | 3.499 | Euro-Immigr | 3.083 | 8.06E-14 |
| CBM58 | 3.411 | Euro-Immigr | 3.040 | 1.71E-13 |
| GH112 | 3.927 | Euro-Immigr | 3.535 | 9.66E-10 |
| GH32 | 4.296 | Euro-Immigr | 3.805 | 2.93E-13 |

**Table S5. Pairwise comparisons of macronutrient intake in participants**

**Male Macronutrient Intake Pairwise Comparisons**

| Macronutrient | Adjusted *P* value |
|---|---|
| **Energy** | |
| Indo-Immigr vs. Euro-Can | 0.0144 [c] |
| **Protein** | |
| Indian vs. Indo-Can | 0.0017[b] |
| Indian vs. Euro-Can | <0.0001[b] |
| Indo-Immigr vs. Indo-Can | 0.0009[b] |
| Indo-Immigr vs. Euro-Can | <0.0001[b] |
| **Fat** | |
| Indian vs. Euro-Can | 0.0175[c] |
| Indo-Immigr vs. Euro-Can | 0.0084[c] |
| **Carbohydrate** | |
| Indian vs. Indo-Immigr | 0.0228 [c] |
| Indian vs. Indo-Can | 0.0177 [c] |
| **Fiber** | |
| Indian vs. Indo-Can | 0.0014 [a] |
| **PUFA** | |
| Indo-Immigr vs. Euro-Can | 0.0140 [a] |
| **Omega-3** | |
| Indo-Immigr vs. Euro-Can | 0.0064 [a] |
| **Omgea-6** | |
| Indo-Immigr vs. Euro-Can | 0.0096 [a] |
| **MUFA** | |
| Indian vs. Euro-Can | 0.0112[a] |
| Indo-Immigr vs. Euro-Can | 0.0154[a] |
| **SFA** | |
| Indian vs. Euro-Can | 0.0312[c] |
| Indo-Immigr vs. Euro-Can | 0.0025[c] |

Multiple Comparisons Test: [a]Dunn’s; [b]Holm-Sidak’s, [c]Tukey’s
Abbreviations: polyunsaturated fatty acid (PUFA); monounsaturated fatty acid (MUFA); saturated fatty acid (SFA)

**Female Macronutrient Intake Pairwise Comparisons**

| Macronutrient | Adjusted *P* value |
|---|---|
| **Protein** | |
| Indian vs. Euro-Can | 0.0008 [a] |
| Indian vs. Euro-Immigr | 0.0143 [a] |
| **Fat** | |
| Indian vs. Euro-Can | 0.0034 [a] |
| Indian vs. Euro-Immigr | 0.0212 [a] |
| **Fiber** | |
| Indian vs. Indo-Can | 0.0487 [a] |
| Indo-Can vs. Euro-Can | 0.0080 [a] |

Multiple Comparisons Test: [a]Dunn’s

**Table S6. Micronutrient intake in males and females**

**Male Participant Absolute Micronutrient Intake**

| **Macronutrient** | **RDA/AMDR** | **Indian**<br>***n = 24*** | **Indo-Immigr**<br>***n = 14*** | **Indo-Can**<br>***n = 11*** | **Euro-Can**<br>***n = 25*** | **Euro-Immigr**<br>***n = 13*** | ***P* value** |
|---|---|---|---|---|---|---|---|
| **Vitamin A, mcg (IQR)** | 600 | 243 (279) | 411 (278) | 365 (332) | 697 (913) | 404 (661) | 0.0007 [a] |
| **Vitamin B1, mg (IQR)** | 1.1 | 1.43 (0.83) | 1.03 (0.67) | 0.73 (0.33) | 0.94 (0.43) | 1.03 (0.68) | 0.0024 [a] |
| **Vitamin B2, mg (IQR)** | 1.1 | 1.01 (0.50) | 0.89 (0.60) | 0.97 (0.29) | 1.50 (0.99) | 1.21 (0.86) | 0.0181* [a] |
| **Vitamin B6, mg (IQR)** | 1.3 | 1.24 (0.90) | 1.07 (0.57) | 0.78 (0.45) | 1.40 (0.90) | 1.64 (1.39) | 0.0276 [a] |
| **Vitamin B12, mcg (IQR)** | 2.4 | 1.24 (1.00) | 1.58 (1.57) | 2.11 (1.91) | 2.11 (1.47) | 3.06 (2.23) | 0.0357* [a] |
| **Vitamin C, mg (IQR)** | 75 | 32.4 (32.4) | 84.3 (74.0) | 42.7 (54.8) | 94.7 (97.5) | 56.2 (130) | 0.0002 [a] |
| **Vitamin D, mcg (IQR)** | 15 | 2.13 (2.25) | 2.53 (2.43) | 1.91 (3.03) | 1.75 (2.65) | 2.39 (3.15) | ns |
| **Zinc, mg (IQR)** | 8 | 6.09 (3.46) | 5.22 (2.83) | 4.28 (2.88) | 6.55 (2.71) | 5.58 (4.13) | 0.0387* [a] |
| **Iron, mg (IQR)** | 18 | 13.6 (9.49) | 11.7 (11.2) | 8.73 (3.32) | 13.0 (3.15) | 13.4 (6.19) | 0.0067 [a] |
| **Folate, mcg (IQR)** | 400 | 365 (223) | 263 (199) | 163 (134) | 267 (210) | 246 (513) | 0.0191 [a] |
| **Magnesium, mg (IQR)** | 310 | 255 (153) | 188 (152) | 130 (47.5) | 263 (152) | 207 (162) | 0.0027 [a] |
| **Calcium, mg (IQR)** | 1000 | 640 (411) | 752 (690) | 631 (271) | 852 (756) | 767 (780) | ns |
| **Sodium, mg (IQR)** | 1500 | 2584 (1809) | 3059 (1949) | 3046 (1394) | 3256 (2027) | 4245 (2663) | ns |

Abbreviations: [a]Kruskal-Wallis, [b] ANOVA. Median and IQR reported.

**Female Participant Absolute Micronutrient Intake**

| Macronutrient | RDA/AMDR | Indian<br>*n = 24* | Indo-Immigr<br>*n = 14* | Indo-Can<br>*n = 11* | Euro-Can<br>*n = 25* | Euro-Immigr<br>*n = 13* | *P* value |
|---|---|---|---|---|---|---|---|
| **Vitamin A, mcg (IQR)** | 600 | 243 (279) | 411 (278) | 365 (332) | 697 (913) | 404 (661) | 0.0007 [a] |
| **Vitamin B1, mg (IQR)** | 1.1 | 1.43 (0.83) | 1.03 (0.67) | 0.73 (0.33) | 0.94 (0.43) | 1.03 (0.68) | 0.0024 [a] |
| **Vitamin B2, mg (IQR)** | 1.1 | 1.01 (0.50) | 0.89 (0.60) | 0.97 (0.29) | 1.50 (0.99) | 1.21 (0.86) | 0.0181* [a] |
| **Vitamin B6, mg (IQR)** | 1.3 | 1.24 (0.90) | 1.07 (0.57) | 0.78 (0.45) | 1.40 (0.90) | 1.64 (1.39) | 0.0276 [a] |
| **Vitamin B12, mcg (IQR)** | 2.4 | 1.24 (1.00) | 1.58 (1.57) | 2.11 (1.91) | 2.11 (1.47) | 3.06 (2.23) | 0.0357* [a] |
| **Vitamin C, mg (IQR)** | 75 | 32.4 (32.4) | 84.3 (74.0) | 42.7 (54.8) | 94.7 (97.5) | 56.2 (130) | 0.0002 [a] |
| **Vitamin D, mcg (IQR)** | 15 | 2.13 (2.25) | 2.53 (2.43) | 1.91 (3.03) | 1.75 (2.65) | 2.39 (3.15) | ns |
| **Zinc, mg (IQR)** | 8 | 6.09 (3.46) | 5.22 (2.83) | 4.28 (2.88) | 6.55 (2.71) | 5.58 (4.13) | 0.0387* [a] |
| **Iron, mg (IQR)** | 18 | 13.6 (9.49) | 11.7 (11.2) | 8.73 (3.32) | 13.0 (3.15) | 13.4 (6.19) | 0.0067 [a] |
| **Folate, mcg (IQR)** | 400 | 365 (223) | 263 (199) | 163 (134) | 267 (210) | 246 (513) | 0.0191 [a] |
| **Magnesium, mg (IQR)** | 310 | 255 (153) | 188 (152) | 130 (47.5) | 263 (152) | 207 (162) | 0.0027 [a] |
| **Calcium, mg (IQR)** | 1000 | 640 (411) | 752 (690) | 631 (271) | 852 (756) | 767 (780) | ns |
| **Sodium, mg (IQR)** | 1500 | 2584 (1809) | 3059 (1949) | 3046 (1394) | 3256 (2027) | 4245 (2663) | ns |

Abbreviations: [a]Kruskal-Wallis. Median and IQR reported.
* Dunn's multiple comparisons revealed no significant differences between groups.

**Male Micronutrient Intake Pairwise Comparisons**

| Macronutrient | Adjusted *P* value |
|---|---|
| **Vitamin A** | |
| Indian vs. Euro-Can | 0.0004 [a] |
| Indo-Immigr vs. Euro-Can | 0.0108 [a] |
| **Vitamin B1** | |
| Indian vs. Indo-Immigr | 0.0021[b] |
| Indo-Immigr vs. Euro-Can | 0.0383[b] |
| **Vitamin B2** | |
| Indian vs. Indo-Can | 0.0038 [b] |
| Indian vs. Euro-Can | <0.0001[b] |
| Indian vs. Euro-Immigr | 0.0172 [b] |
| Indo-Immigr vs. Indo-Can | 0.0076 [b] |
| Indo-Immigr vs. Euro-Can | <0.0001[b] |
| Indo-Immigr vs. Euro-Immigr | 0.0373 [b] |
| **Vitamin B12** | |
| Indian vs. Euro-Can | 0.0099 [a] |
| Indo-Immigr vs. Euro-Can | 0.0194 [a] |
| **Vitamin D** | |
| Indian vs. Euro-Can | 0.0112 [a] |
| **Zinc** | |
| Indian vs. Indo-Immigr | <0.0001 [a] |
| Indian vs. Indo-Can | 0.0022 [a] |
| Indian vs. Euro-Can | 0.0134 [a] |
| Indian vs. Euro-Immigr | <0.0001 [a] |
| **Iron** | |
| Indian vs. Indo-Immigr | 0.0048 [b] |
| **Folate** | |
| Indian vs. Indo-Immigr | 0.0047 [b] |
| Indian vs. Euro-Immigr | 0.0304 [b] |
| **Magnesium** | |
| Indo-Immigr vs. Euro-Can | 0.0173 [b] |

| | |
|---|---|
| **Calcium** | |
| Indian vs. Euro-Can | 0.0304 [a] |

Multiple Comparisons Test: [a]Dunn’s, [b]Tukey’s

**Female Micronutrient Intake Pairwise Comparisons**

| **Macronutrient** | **Adjusted *P* value** |
|---|---|
| **Vitamin A** | |
| Indian vs. Euro-Can | 0.0001[a] |
| **Vitamin B1** | |
| Indian vs. Indo-Can | 0.0007[a] |
| **Vitamin B6** | |
| Indo-Can vs. Euro-Immigr | 0.0268 [a] |
| **Vitamin C** | |
| Indian vs. Indo-Immigr | 0.0463 [a] |
| Indo-Immigr vs. Euro-Can | 0.0001 [a] |
| **Iron** | |
| Indian vs. Indo-Can | 0.0284 [a] |
| Indo-Can vs. Euro-Can | 0.0148 [a] |
| Indo-Can vs. Euro-Immigr | 0.0070 [a] |
| **Folate** | |
| Indian vs. Indo-Can | 0.0062 [a] |
| **Magnesium** | |
| Indian vs. Indo-Can | 0.0070 [a] |
| Indo-Can vs. Euro-Can | 0.0026 [a] |
| Indo-Can vs. Euro-Immigr | 0.0496 [a] |

Multiple Comparisons Test: [a]Dunn’s
